# A phase-separated biomolecular condensate nucleates polymerization of the tubulin homolog FtsZ to spatiotemporally regulate bacterial cell division

**DOI:** 10.1101/2022.09.12.507586

**Authors:** Beatrice Ramm, Dominik Schumacher, Andrea Harms, Tamara Heermann, Philipp Klos, Franziska Müller, Petra Schwille, Lotte Søgaard-Andersen

## Abstract

Cell division is spatiotemporally precisely regulated, but the underlying mechanisms are incompletely understood. In the social, predatory bacterium *Myxococcus xanthus*, the PomX/PomY/PomZ proteins form a single large megadalton-sized complex that directly positions and stimulates cytokinetic ring formation by the tubulin homolog FtsZ. Here, we studied the structure and mechanism of this complex *in vitro* and *in vivo.* We demonstrate that PomY forms liquid-like biomolecular condensates by phase separation, while PomX self-assembles into filaments generating a single large cellular structure. The PomX structure enriches PomY, thereby guaranteeing the formation of precisely one PomY condensate per cell through surface-assisted condensation. *In vitro*, PomY condensates selectively enrich FtsZ and nucleate GTP-dependent FtsZ polymerization, suggesting a novel cell division site positioning mechanism in which the single PomY condensate enriches FtsZ to guide FtsZ-ring formation and division. PomY-nucleated FtsZ polymerization shares features with microtubule nucleation by biomolecular condensates in eukaryotes, supporting this mechanism’s ancient origin.

## Introduction

Although they generally lack organelles, bacterial cells are spatiotemporally highly organized with proteins localizing dynamically to distinct subcellular locations to spatially restrict their activities even (Surovtsev and Jacobs-Wagner, 2018; Treuner-Lange and Søgaard-Andersen, 2014). However, our understanding of how this spatiotemporal organization is accomplished is incomplete. Recently, biomolecular condensates formed by liquid-liquid phase separation (LLPS) have emerged as an important mechanism to spatially organize intracellular processes in eukaryotic cells (Alberti and Hyman, 2021; Banani et al., 2017; Lyon et al., 2021; Shin and Brangwynne, 2017), while they are only beginning to be identified and explored in bacteria (Azaldegui et al., 2021).

The formation of biomolecular condensates in bulk solution involves the concentration-dependent, switch-like demixing of proteins from solution into high-density condensates that coexist with a remaining dilute protein phase above a critical saturation concentration (*C*_sat_) (Alberti and Hyman, 2021; Banani et al., 2017; Lyon et al., 2021; Shin and Brangwynne, 2017). Alternatively, condensates can be nucleated through surface-assisted condensation (Banjade and Rosen, 2014; Hernandez-Vega et al., 2017; Morin et al., 2022; Snead et al., 2022; Tan et al., 2019). This might arise from a phenomenon termed prewetting (Cahn, 1977; Nakanishi and Fisher, 1982), in which the binding of a protein to a surface results in a local enrichment thereby initiating condensate formation on the surface. As the bulk concentration is below *C*_sat_, condensate formation is restricted to the surface, thereby providing a means of spatiotemporal regulation. If the bulk concentration is above *C*_sat_, condensates formed in the bulk can also associate with the surface to wet it (Gouveia et al., 2022; Wiegand and Hyman, 2020).

For all condensates, the interface between the two phases forms a boundary that serves as a selective barrier for some molecules but not for others, resulting in selective enrichment of so-called client proteins and/or RNA molecules (Alberti and Hyman, 2021; Banani et al., 2017; Lyon et al., 2021; Shin and Brangwynne, 2017). In this way, biomolecular condensates can enhance chemical reactions, sequester molecules, or act as hubs to nucleate microtubule or actin polymerization (Banjade and Rosen, 2014; Graham et al., 2022; Hernandez-Vega et al., 2017; King and Petry, 2020; Li et al., 2012; Woodruff et al., 2017; Yang et al., 2022).

Protein condensates form by cumulative, specific, transient, low-affinity interactions between multivalent proteins that may self-interact via homotypic interactions or interact with other proteins via heterotypic interactions using folded domains, intrinsically disordered regions (IDRs), and/or repetitive protein motifs in low-complexity regions (Banani et al., 2017). Above the *C*_sat_, the multivalent interactions give rise to large non-stoichiometric networks of interacting proteins that lead to condensate formation (Alberti and Hyman, 2021; Banani et al., 2017; Lyon et al., 2021; Shin and Brangwynne, 2017). Many condensates have liquid-like properties and are of spherical shapes due to the surface tension acting to minimize the surface area (Alberti and Hyman, 2021; Banani et al., 2017; Lyon et al., 2021; Shin and Brangwynne, 2017). Such liquid-like condensates can be highly dynamic and exchange molecules with the dilute phase, undergo internal reorganization, fusion, fission, and/or disintegration (Alberti and Hyman, 2021; Banani et al., 2017; Lyon et al., 2021; Shin and Brangwynne, 2017).

Positioning of the cell division site in bacteria is spatiotemporally precisely regulated. Bacterial cell division depends on the localization of the tubulin-homolog FtsZ at the incipient division site to form, in a GTP-dependent manner, the so-called Z-ring (Lutkenhaus et al., 2012; McQuillen and Xiao, 2020), a ring-like structure of short treadmilling FtsZ filaments (Bisson-Filho et al., 2017). Subsequently, the Z-ring directly or indirectly recruits other proteins that help to execute cytokinesis (Lutkenhaus et al., 2012; McQuillen and Xiao, 2020). The systems that spatiotemporally control cell division accomplish their function by directly interacting with FtsZ to modulate its GTP-dependent polymerization. Negative regulation systems such as the MinC/MinD/MinE system in *Escherichia coli* and the MipZ/ParB system in *Caulobacter crescentus* position an FtsZ inhibitor at the cell poles, whereby Z-ring formation is restricted to midcell (Kiekebusch et al., 2012; Ramm et al., 2019). By contrast, MapZ in the pathogen *Streptococcus pneumoniae* (Fleurie et al., 2014; Holeckova et al., 2014) and the PomX/PomY/PomZ proteins in the rod-shaped cells of the social, predatory bacterium *Myxococcus xanthus* localize at midcell to directly guide and promote Z-ring formation and cell division between two segregated chromosomes (Harms et al., 2013; Schumacher et al., 2017; Treuner-Lange et al., 2013).

The PomX/PomY/PomZ system is a representative of a large group of systems that include the MinC/MinD/MinE system for cell division placement and the ParA/ParB system for chromosome and plasmid segregation, and in which a ParA/MinD-type ATPase together with its cognate ATPase activating protein (AAP) positions macromolecular structures in bacteria (Lutkenhaus, 2012). In the PomX/PomY/PomZ system, the AAPs PomX and PomY form a single large cytoplasmic complex that is recruited to the nucleoid by the ATP-bound dimeric ParA/MinD-type ATPase PomZ (Schumacher et al., 2017; Treuner-Lange et al., 2013). PomX, PomY and PomZ are present in multiple copies in this complex, which has an average size of ~15MDa and is visible by widefield fluorescence microscopy with the three proteins colocalizing (Schumacher et al., 2017). After cell division, the PomX/PomY/PomZ complex is bound to the nucleoid close to the new cell pole. Subsequently, it translocates by biased random motion to midcell, where it switches to constrained motion and stimulates Z-ring formation by an unknown mechanism (Schumacher et al., 2017). Upon division, the PomX/PomY/PomZ complex was suggested to undergo fission with both daughters “receiving” part of the complex (Schumacher et al., 2017). The three Pom proteins interact in all pairwise combinations and have distinct functions (Schumacher et al., 2017; Schumacher et al., 2021): As shown by negative stain transmission electron microscopy (TEM), PomX self-assembles to form filaments *in vitro. In vivo*, PomX assembles into a single cellular cluster independently of PomY and PomZ and stimulates the assembly of the PomY and PomZ clusters by direct interaction to generate the PomX/PomY/PomZ complex. PomY interacts directly with FtsZ and is essential for its recruitment to the incipient division site. PomZ interacts directly with PomX and PomY and associates the PomX/PomY complex with the nucleoid. By stimulating ATP hydrolysis by DNA-bound PomZ, PomX and PomY promote cluster translocation.

While PomX/PomY/PomZ cluster translocation is well-understood based on experiments and theory (Bergeler and Frey, 2018; Kober et al., 2019; Schumacher et al., 2017), the structure of the PomX/PomY/PomZ complex and its function in Z-ring formation remain unclear. Here, by focusing on PomX and PomY and combining *in vivo* and *in vitro* approaches, we show that PomY forms liquid-like biomolecular condensates by phase separation. These condensates selectively enrich FtsZ and nucleate GTP-dependent FtsZ polymerization. The filamentous PomX structure serves as a scaffold to locally enrich PomY, thereby nucleating phase separation by PomY via surface-assisted condensation and ensuring the formation of precisely one PomY condensate per cell that guides Z-ring formation and cell division.

## Results

### PomX and PomY clusters are structurally flexible, non-stoichiometric and scale in size with cell size

To understand the structure and function of the PomX/PomY/PomZ complex *in vivo*, we focused on PomX and PomY because they form a single cytoplasmic complex independently of PomZ that is able to stimulate Z-ring formation, while the PomZ ATPase is important for translocation of this complex (Schumacher et al., 2017). In strains expressing active mCherry (mCh) fusions, i.e. mCh-PomX and PomY-mCh, at native levels, we observed that cluster dimensions were above the diffraction limit of high-resolution structured illumination microscopy (SIM) (Fig. 1AB; Fig. S1A-C). Individual cells generally contained a single cluster, but smaller cells sometimes lacked a visible cluster (Fig. S1D). For both proteins, cluster shape varied from spherical to spheroidal (Fig. 1AB). Generally, mCh-PomX clusters were more elongated than PomY-mCh clusters while the short axes were comparable (Fig. 1B). Accordingly, the mean aspect ratio of mCh-PomX clusters was significantly higher than that of PomY-mCh clusters (Fig. 1B). Interestingly, for both proteins, the spherical clusters tended to be present in shorter cells while longer cells contained the larger spheroidal clusters (Fig. 1C). Because the short cluster axes were independent of cell length (Fig. 1C), we conclude that cluster shape varies from spherical to spheroidal and cluster size scales with cell size.

**Fig. 1.**
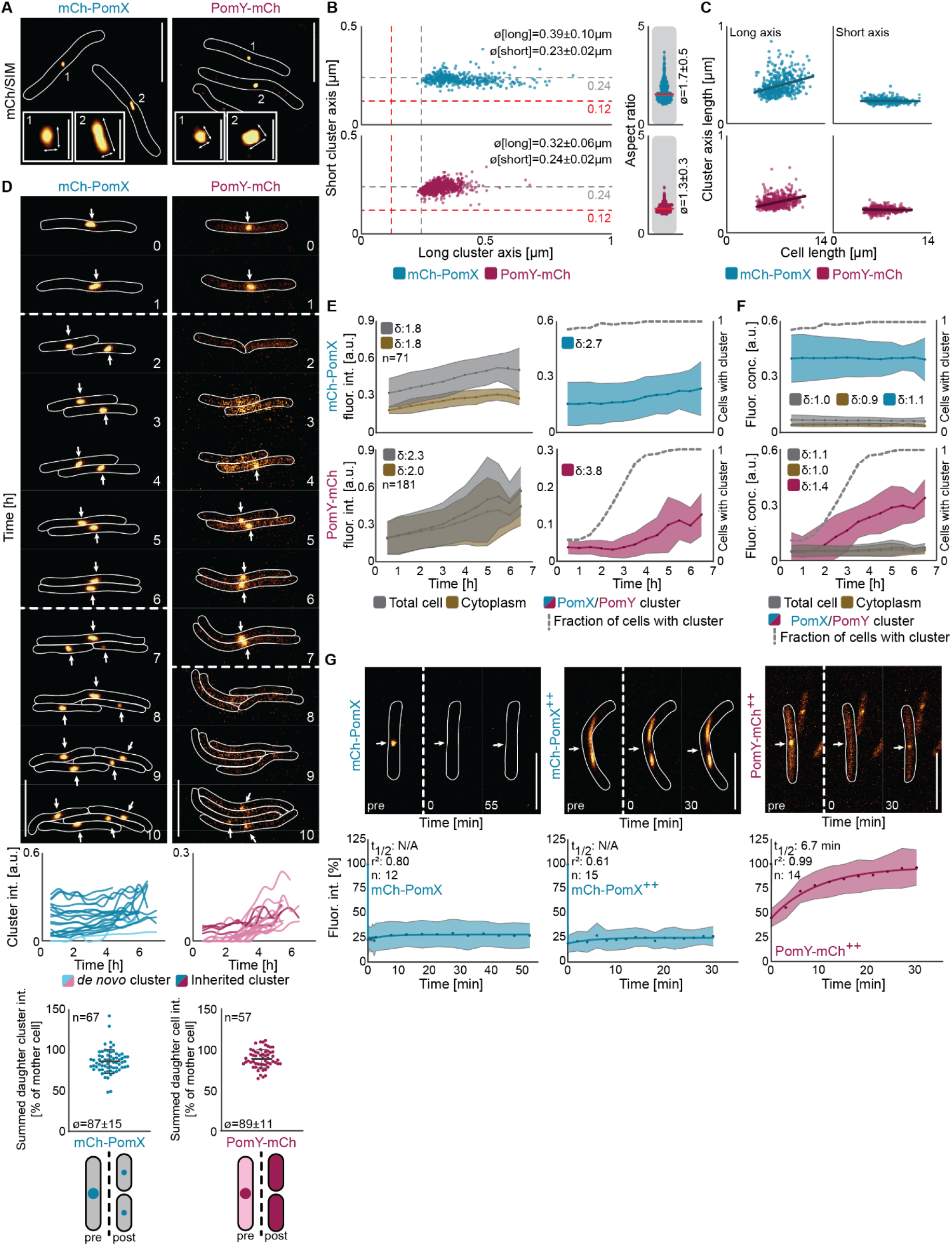
PomX and PomY clusters are structurally flexible, non-stoichiometric and scale in size with cell size. A. SIM images of cells expressing mCh-PomX or PomY-mCh at native levels. Numbers indicate numbering of the insets. Arrows indicate long and short cluster axes used for quantification. Cell outlines are shown as white lines. Scale bars, 5μm and 0.5μm in insets. B. Quantification of long and short cluster axes and cluster aspect ratios based on SIM images. Red and grey stippled lines in the left panels indicate the theoretical resolution of SIM and epifluorescence microscopy, respectively; red solid lines in the right panels indicate the median. n >400 clusters in at least two independent replicates. ø indicates mean±standard deviation (STDEV). C. Quantification of long and short cluster axes as function of cell length based on SIM images. Dark-colored lines indicate linear regression. The same data as in B were used for the analysis. D. Epifluorescence time-lapse microscopy of cells expressing mCh-PomX or PomY-mCh at native levels. Stippled lines indicate cell divisions. Arrows point to mCh-PomX and PomY-mCh clusters. Scale bars, 5 μm. Middle panels, quantification of mCh-PomX and PomY-mCh cluster intensity in 25 representative cells from birth to division. Dark-, and light-colored lines show intensity of clusters that were inherited from the mother cell and *de novo* synthesized, respectively. Lower panels, (left) quantification of mCh-PomX cluster fluorescence intensity pre- and post-division in mother cells and their corresponding daughters; (right) quantification of total cellular PomY-mCh fluorescence intensity pre- and post-division in mother cells and their corresponding daughters. n, number of divisions analyzed; error bars, mean±STDEV. E. Fluorescence intensity of cells expressing mCh-PomX or PomY-mCh at native levels over the cell cycle. Colored dotted lines indicate the mean and light-colored areas STDEV. δ values indicate the fold increase between birth and division. n, number of cells analyzed. F. Fluorescence concentrations of cells expressing mCh-PomX or PomY-mCh at native levels over the cell cycle. Same cells as in E were used. G. Fluorescence recovery after photobleaching (FRAP) experiments. Cluster fluorescence intensity in a region of interest (ROI) of 0.52μm in diameter before bleaching was set to 100%. Colored dots indicate the mean and light-colored areas STDEV. Dark lines show the recovery fitted to a single exponential equation. ++ indicates that the proteins were moderately overexpressed from the *pilA* promoter. Stippled lines indicate bleaching events marked by white arrows. n, number of bleaching events. Scale bars, 5μm. See also Figure S1.

We next examined the size-scaling of the mCh-PomX and PomY-mCh clusters by quantitative wide-field time-lapse microscopy and using fluorescence intensity as a metric for cluster size. All cells had mCh-PomX and PomY-mCh clusters at midcell immediately before cell division (Fig. 1D). For mCh-PomX, most daughters contained a cluster after cell division that, generally, was smaller than the one of the mother (Fig. 1D). Because the summed fluorescence intensities of the daughter clusters added up to 87±15% of the mother’s cluster (Fig. 1D), we conclude that the mCh-PomX clusters undergo fission during cell division. Sometimes the fission process was asymmetric with two daughters receiving clusters of unequal sizes (Fig. 1D). Occasionally, a daughter did not “receive” a mCh-PomX cluster; such cells subsequently formed a cluster *de novo* (Fig. 1D). mCh-PomX clusters essentially doubled in size over the cell cycle as did total cellular fluorescence, and cytoplasmic fluorescence (Fig. 1DE). By normalizing fluorescence intensities to the respective areas to obtain a metric for protein concentration (referred to as fluorescence concentration), we found that the fluorescence concentration of mCh-PomX in the cell, the cytoplasm, and the cluster remained constant during the cell cycle (Fig. 1F). Remarkably, most PomY-mCh clusters disappeared during cell division resulting in daughters with only diffuse PomY-mCh signals that added up to 89±11% of the mother’s total cellular fluorescence (Fig. 1D), supporting that PomY-mCh in clusters is not proteolytically degraded during division, but rather the clusters undergo disintegration during division. Within hours of a cell division, a single weak cluster emerged *de novo* and grew in size before the subsequent cell division, resulting in cycles of cluster growth and disintegration (Fig. 1D). For PomY-mCh, total cellular fluorescence and cytoplasmic fluorescence also essentially doubled over the cell cycle, while cluster fluorescence intensity (and therefore cluster size) increased ~4-fold; this increase represents a minimum estimate because it is calculated from the first appearance of a cluster until the subsequent cell division (Fig. 1E). Moreover, the cellular and cytoplasmic PomY-mCh fluorescence concentrations remained constant over the cell cycle, while cluster fluorescence concentration increased ~40% (Fig. 1F); as above, this represents a minimum estimate.

In snapshots of 1000s of cells and using cell length as a proxy for the cell cycle stage, we confirmed the results from the time-lapse microscopy for total cellular, cytoplasmic, and cluster fluorescence as well as for fluorescence concentrations (Fig. S1EF). Based on these experiments, we calculated a mean enrichment factor in a cluster relative to the cytoplasm of 12.1 for mCh-PomX and 4.1 for PomY-mCh. Based on the estimated PomX and PomY concentrations in wild-type (WT) cells (Schumacher et al., 2017), the mean mCh-PomX concentration in a cluster and the cytoplasm is 1.0μM and 0.08μM, respectively and the corresponding PomY-mCh concentrations 1.4μM and 0.3μM.

Because the PomX and PomY clusters grow in size with time, we asked whether the proteins dynamically exchange with the cytoplasm. In fluorescence recovery after photobleaching (FRAP) experiments, neither mCh-PomX clusters at native protein levels nor when moderately overexpressed showed recovery (Fig. 1G). PomY-mCh signals were too weak for FRAP analyses at native protein levels; however, when PomY-mCh was moderately overexpressed, PomY-mCh in clusters dynamically exchanged with the cytoplasm with a half-maximal recovery time (t1/2) of 6-7 min (Fig. 1G).

In summary, for both proteins cluster shape varies from spherical to spheroidal and cluster size scales directly with cell size despite the constant cytoplasmic and cellular concentrations. PomX clusters undergo fission during cell division and grow in size, while PomY clusters disintegrate during division and then reform *de novo* followed by growth. Moreover, PomY in clusters dynamically exchanges molecules with the cytoplasm, while PomX does not. We conclude that the PomX/PomY complex is structurally and compositionally highly flexible, neither the PomX nor the PomY part of this complex has a fixed stoichiometry, and the combined complex lacks a fixed PomX/PomY stoichiometry. This is unlike structures with a fixed stoichiometry, e.g. the ribosome, that are held together by precise, stereospecific interactions and do not exhibit size scaling. Importantly, spherical to spheroidal cellular clusters, non-stoichiometric complexes, direct size scaling with cell size in combination with a constant cytoplasmic concentration (Brangwynne, 2013; Weber and Brangwynne, 2015), dynamic exchange of components, as well as cluster fission and disintegration are defining features of biomolecular condensates formed by LLPS, suggesting that LLPS might also be involved in the formation of the PomX/PomY complex.

### PomX forms filamentous structures and PomY forms biomolecular condensates *in vitro*

To determine whether PomX and PomY share characteristics with proteins that undergo LLPS, we analyzed the two proteins bioinformatically (Fig. 2A). PomX consists of an N-terminal, disordered region (PomX^IDR^), which includes the conserved N-terminal 22 residues (PomX^NPEP^) that are sufficient to stimulate ATP hydrolysis by PomZ (Schumacher et al., 2021), as well as a C-terminal α-helical region predicted to form a coiled-coil (PomX^CC^). This architecture is conserved in PomX orthologs (Schumacher et al., 2021). PomX^CC^ is required and sufficient for PomX polymerization and also interacts with PomY; both parts are required for PomX function (Schumacher et al., 2021). PomY comprises an N-terminal α-helical region predicted to form a coiled-coil (PomY^CC^), followed by a region with six HEAT-repeats (PomY^HEAT^), and a C-terminal mostly disordered region (PomY^IDR^). This architecture is conserved in PomY orthologs (Fig. S2A). All five parts of the two proteins have a biased amino acid composition and also contain a large number of charged residues (Fig. 2A) resulting in a pI of 4.9 and 9.3 for PomX and PomY, respectively. Thus, both proteins possess the hallmarks of proteins that undergo LLPS, including multiple domains that might engage in protein-protein interactions, IDRs, and low sequence complexity.

**Fig. 2.**
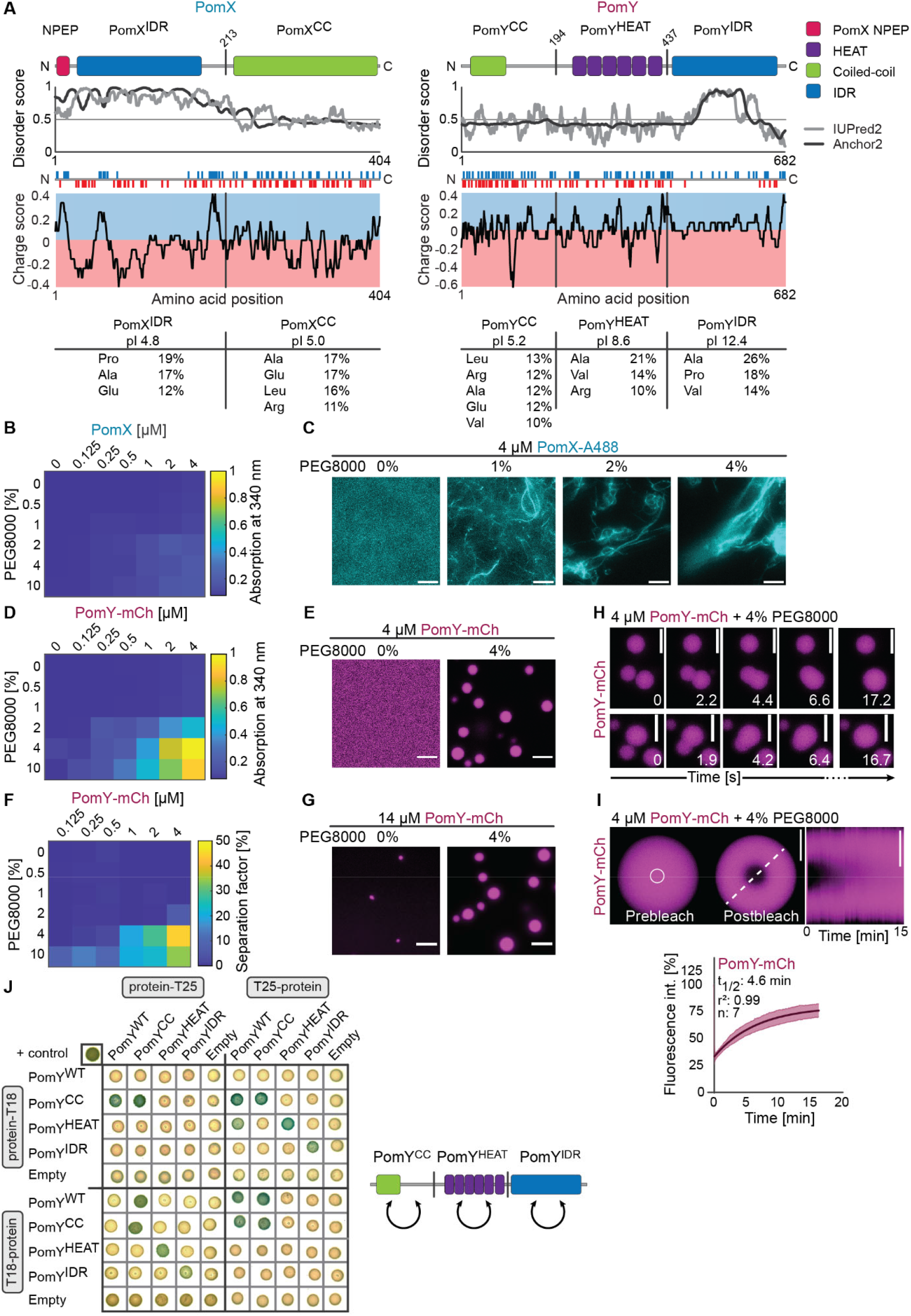
PomX forms filamentous structures and PomY biomolecular condensates *in vitro*. A. Domain architecture of PomX and PomY. PomX domain truncations used in (Schumacher et al., 2021) and PomY truncations as used here are indicated by vertical lines. PomX^IDR^ extends from residue 1-213, PomX^CC^ from 214-404, PomY^CC^ from 1-194, PomY^HEAT^ from 195-437, and PomY^IDR^ from 438-682. In red and blue, positively (Arg, Lys, His) and negatively charged (Glu, Asp) residues are indicated. Charge score was plotted using a sliding window of six residues. Blue and red indicate positively and negatively charge regions, respectively. Amino acid composition is shown for individual domains for amino acids that make up >10% of residues. B. Turbidity measurements of PomX for increasing protein and PEG8000 concentrations. The heat map displays average absorption at 340 nm for n=3 replicates. C. PomX forms filaments that are increasingly bundled with increasing crowder concentrations. Representative images of 4μM PomX-A488 in the presence of increasing concentrations of PEG8000. Scale bars, 50μm. D-G. PomY phase separates in a protein and crowding-agent dependent manner. D. Turbidity measurements of PomY-mCh with increasing protein and PEG8000 concentrations. Heat map displays average absorption at 340 nm for n=6 replicates. E. Representative images of 4μM PomY-mCh in absence and presence of 4% PEG8000. Scale bars, 5μm. F. Heat map of PomY-mCh phase separation with increasing protein and crowder concentrations. Values are the average separation factor in percent calculated from the maximum intensity projection of confocal z-stacks. Data from two independent experiments with n=8 analyzed images per condition. G. Representative images of 14μM PomY in the absence and presence of 4% PEG8000. Scale bars, 5μm. H. Time-series of PomY-mCh condensates undergoing fusion (4μM PomY-mCh, 4% PEG8000). Scale bars, 5μm. I. Partial FRAP of PomY-mCh condensates (4μM PomY-mCh, 4% PEG8000). Representative images of a PomY-mCh condensate before and after bleaching (white circle indicates ROI). Kymograph of the fluorescence recovery was created along the white stippled line. In the diagram below, fluorescence intensity in the ROI was set to 100% before bleaching; dark magenta line indicates the mean recovery, fitted to a single exponential equation with the light-colored area showing the STDEV from n=7 bleaching events. Scale bars, 5μm. J. BACTH analysis of PomY and PomY domains. The indicated PomY domains (see A) were fused to the N- and the C-terminus of T18 and T25 as indicated. Blue and white colony colors indicate an interaction and no interaction, respectively. Positive control in upper left. The image shows representative results of three independent experiments. Right panel, schematic of the observed interactions indicated by arrows. See also Figures S2 and S3 and Movie 1.

To determine whether PomX and/or PomY undergo LLPS with condensate formation, we used an *in vitro* assay with purified His_6_-tagged full-length proteins without (referred to as PomX, PomY) or with an mCh-tag (referred to as mCh-PomX, PomY-mCh), as well as a His_6_-tagged PomX variant with a Cys-tag for labeling with the Alexa488-maleimide fluorophore (PomX-A488) (Fig. S2B). These proteins were used to collect phase diagrams based on turbidity measurements and fluorescence microscopy with increasing concentrations of purified protein close to the cellular concentrations (see above) and the crowding agent polyethylene glycol 8000 (PEG8000) to mimic the crowded intracellular environment (Kuznetsova et al., 2014).

The turbidity of PomX and mCh-PomX solutions was similar for all protein and crowding agent concentrations (Fig. 2B, Fig. S2C). In fluorescence microscopy, mCh-PomX and PomX-A488 were inhomogeneously distributed and formed filamentous structures with increasing protein and crowding agent concentrations (Fig. 2C, Fig. S2DE). These observations are in agreement with previous TEM analyses in which PomX forms regular filaments with a width of ~10 nm in the absence of PEG8000 (Schumacher et al., 2017; Schumacher et al., 2021). Spherical mCh-PomX and PomX-A488 condensates were not detected under any condition. We conclude that PomX forms filaments with sizes below the diffraction limit that become increasingly bundled as the protein and PEG8000 concentrations are increased. Together with the observation that PomX clusters grow but do not exchange molecules with the cytoplasm *in vivo*, these observations suggest that PomX does not undergo LLPS, but forms protein filaments, which *in vivo* assemble to form the single PomX cluster.

By contrast, the turbidity of PomY (Fig. S3A) and PomY-mCh (Fig. 2D) solutions increased in a protein and PEG8000 concentration-dependent manner. By microscopy, both proteins formed micrometer-sized, spherical condensates indicative of LLPS under the conditions with increased turbidity (Fig. 2E; S3BC), suggesting that the dense phase has liquid-like properties with the surface tension seeking to minimize the surface area. As a measure for the phase separation behavior of PomY-mCh, we quantified the total amount of protein that segregated into the dense phase by determining the separation factor defined as the integrated fluorescence intensity of the condensates over the total integrated fluorescence intensity in the images. In the resulting phase diagram, conditions with a high separation factor coincided with those that also showed increased turbidity (Fig. 2F). At these PomY and PomY-mCh concentrations, which are close to the cellular concentration, the proteins only formed condensates in the presence of PEG8000. Importantly, at a concentration of 14μM, which is far higher than the estimated cellular concentration, PomY-mCh also formed small spherical condensates in the absence of PEG8000 (Fig. 2G). In addition to their spherical shape, PomY-mCh condensates also displayed other liquid-like behaviors with fusion of condensates followed by relaxation into larger spherical condensates within seconds to minimize surface area (Fig. 2H; Movie 1). Likewise, PomY-mCh condensates were dynamic, as shown by FRAP with recovery after bleaching on a minute timescale (t_1/2_ 4-5 min), similarly to PomY-mCh clusters *in vivo* (Fig. 2I, cf. Fig. 1G). In conclusion, (1) the protein and crowding agent concentration-dependent condensate formation, (2) their liquid-like properties as evidenced by their sphericity, fusion/relaxation events driven by surface tension, and the dynamic exchange of molecules, and (3) the bioinformatic analysis, directly support that PomY in the absence of crowding agent undergoes LLPS at very high concentrations and that *C*_sat_ for LLPS by PomY is lowered in the presence of crowding agent *in vitro.*

Phase separation of PomY-mCh was not specific to the crowding agent used (Fig. S3D), and occurred with various buffers in the physiological pH range of 6.5-8.0 (Fig. S3E). It was independent of MgCl2 and decreased with increasing KCl concentration (Fig. S3F), as well as with increasing concentrations of the aliphatic alcohol 1,6-hexanediol, which perturbs hydrophobic interactions (Ribbeck and Görlich, 2002), (Fig. S3G), suggesting that PomY condensate formation depends on electrostatic as well as hydrophobic interactions.

The phase separation behavior of PomY *in vitro* suggests that PomY self-interacts. Accordingly, using a bacterial adenylate cyclase two-hybrid (BACTH) assay with truncated PomY variants, we observed that full-length PomY as well as the three individual PomY domains, i.e. PomY^CC^, PomY^Heat^ and PomY^IDR^ (Fig. 2A), self-interact (Fig. 2J). We conclude that (1) PomY is a multivalent protein that self-interacts using multivalent homotypic interactions, and (2) PomY phase separation with condensate formation is driven by homotypic electrostatic and hydrophobic interactions.

### PomY condensates wet PomX filaments *in vitro*

*In vivo* PomX cluster formation is independent of PomY, but PomY depends on PomX for cluster formation at native protein levels (Schumacher et al., 2017). Prompted by these observations, we asked whether PomX could stimulate phase separation by PomY *in vitro*. At low PEG8000 concentrations (0-1%), where PomX-A488 alone forms filaments and PomY-mCh alone is homogeneously distributed, the two proteins colocalized in filamentous structures with PomY-mCh coating and bundling PomX filaments (Fig. 3A); this is in agreement with previous TEM analyses demonstrating that PomY coats and bundles PomX filaments (Schumacher et al., 2017). At higher PEG8000 concentrations (2-10%), where PomX-A488 alone forms bundled filaments and PomY-mCh alone phase separates, PomY-mCh not only coated and bundled the PomX-A488 filaments but also formed condensates that often localized on the PomX-A488 bundles (Fig. 3A). Qualitatively these PomY condensates tended to be less spherical than those formed by PomY alone. Moreover, in time-lapse microscopy, we observed that PomY-mCh condensates formed in solution associated with and wetted the filamentous PomX-A488 structures (Fig. 3B). These PomY-mCh condensates also underwent fusion on the PomX-A488 structures, becoming more elongated over time (Fig. 3B; Movie 2). We conclude that at bulk PomY concentrations below *C*_sat_ for LLPS PomX filaments enrich PomY resulting in a high local concentration of PomY on the PomX structure. At higher crowder concentrations, i.e. above the bulk *C*_sat_, PomY condensates also form in solution and associate with and wet the PomX filaments.

**Fig. 3.**
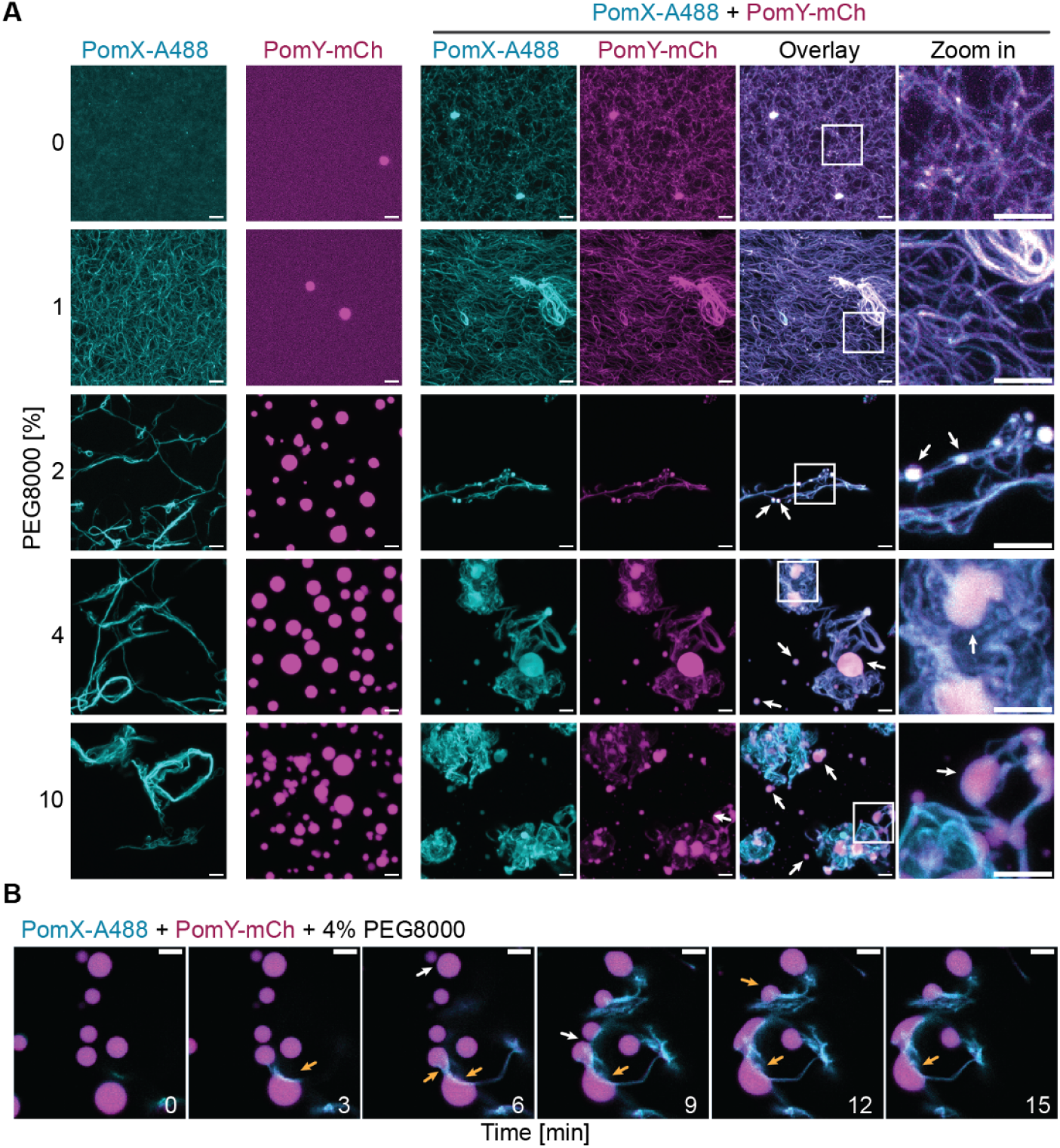
PomY condensates wet PomX filaments *in vitro*. A. Representative maximum intensity projections of confocal z-stacks of 4μM PomX-A488 and 4μM PomY-mCh alone and in combination in the presence of increasing PEG8000 concentrations after 1h of incubation. White frames indicate regions in the zoomed images. White arrows indicate PomY-mCh condensates. Scale bars, 5μm. B. Time-series of PomY-mCh condensates wetting PomX-A488 filaments and deforming upon contact (4μM PomY-mCh, 4μM PomX-A488, 4% PEG8000). White arrows indicate fusion events of PomY-mCh condensates and orange arrows indicate PomY-mCh condensates that deform upon interaction with PomX-A488 filaments. Scale bars, 5μm. See also Movie 2.

### PomY forms functional biomolecular condensates in a concentration-dependent manner *in vivo* in the absence of PomX

Based on the *in vivo* observations that PomY cluster formation depends on PomX and that phase separation might be involved in PomY cluster formation, we hypothesized that the PomY concentration in WT cells is below *C*_sat_ for LLPS. Therefore, in the absence of PomX, PomY is homogenously distributed. This hypothesis predicts that if the cellular PomY concentration is increased sufficiently to exceed *C*_sat_, then PomY should be able to undergo LLPS with condensate formation independently of PomX. To test this hypothesis, we expressed *pomY-mCh* from a vanillate-inducible promoter in strains with or without PomX. At the highly elevated PomY-mCh concentration at 9h of induction (Fig. S4AB) and in the presence of PomX, PomY-mCh formed a single cluster in most cells, but 12% of cells contained multiple clusters (Fig. 4AB). Intriguingly, at this elevated concentration, PomY-mCh formed one or more clusters in the absence of PomX at random positions along the cell length in 41% of cells, while mCh alone was homogenously distributed (Fig. 4AB, Fig. S4CD). These PomY-mCh clusters were highly dynamic, grew in intensity, and were stable over long periods or rapidly disintegrated (Fig. 4C), indicating a dynamic rearrangement of PomY molecules between clusters and the cytoplasm and that these clusters are biomolecular condensates.

**Fig. 4.**
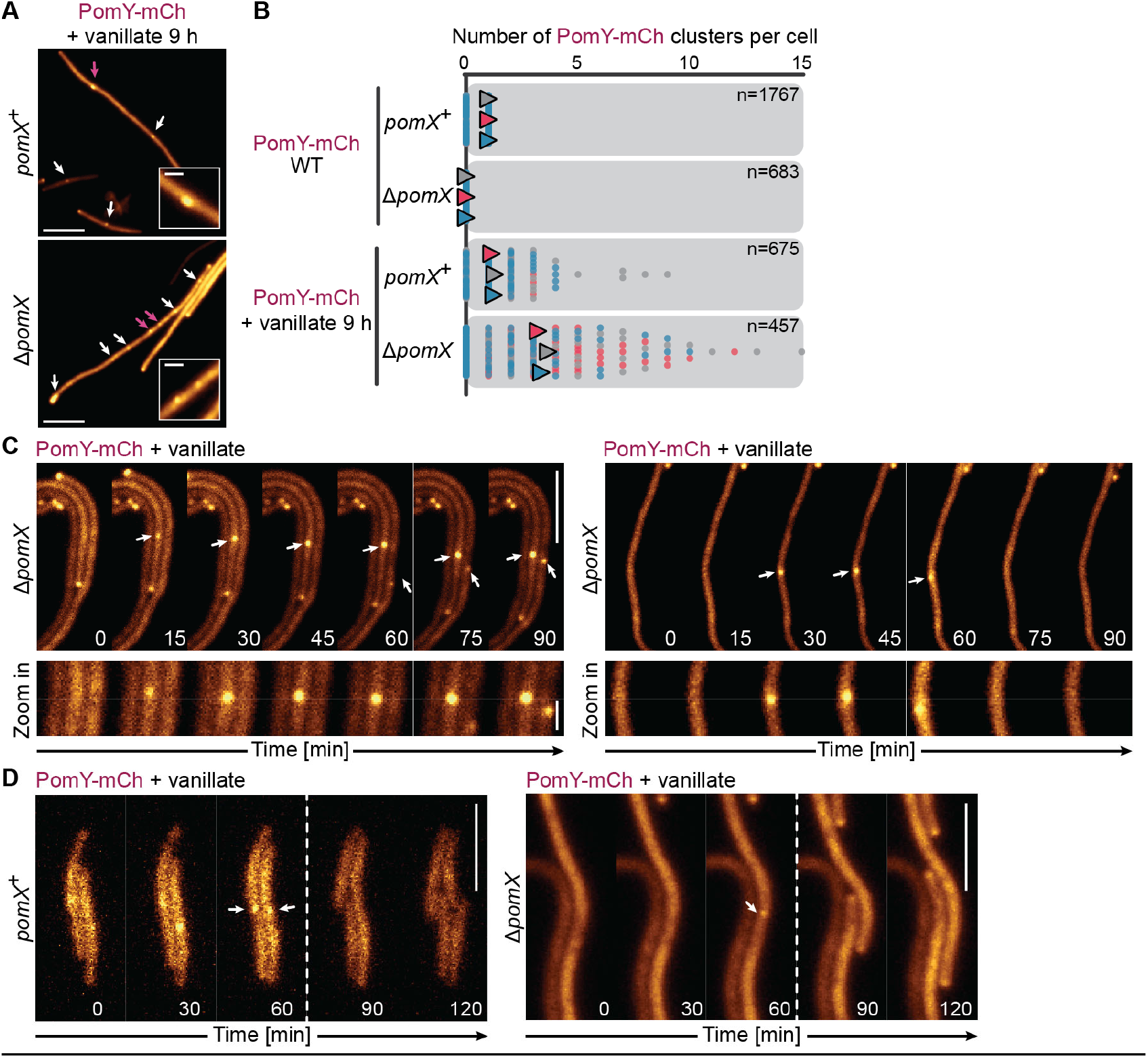
PomY forms functional condensates in a concentration-dependent manner *in vivo* independently of PomX. A. Fluorescence microscopy of cells with a highly elevated PomY-mCh concentration after 9h of induction with 1mM vanillate. White arrows, cytoplasmic PomY-mCh clusters; magenta arrows, clusters shown in insets. Scale bars, 5μm and 0.5μm in insets. B. Quantification of the number of PomY-mCh clusters in cells in A and in cells accumulating PomY-mCh at native levels. Data from three replicates are shown in different colors and with triangles indicating the mean. For calculating the mean, only cells with clusters were included. N, number of cells analyzed. C. Fluorescence time-lapse microscopy of cells with a highly elevated PomY-mCh concentration. Cells grown in suspension without vanillate were transferred to agarose pads containing 1mM vanillate 32°C. Images show cells after ~10h of induction. White arrows indicate cell regions with clusters shown in the zoomed images. Scale bars, 5μm and 0.5μm in insets. D. Fluorescence time-lapse microscopy of cells with a highly elevated PomY-mCh concentration during division. Cells were treated as in C. White arrows indicate clusters immediately before cell division. Stippled lines indicate cell division events. Scale bars, 5μm. See also Figure S4.

To investigate the functionality of the “pure” PomY-mCh condensates formed in the absence of PomX, we analyzed the localization of cell division constrictions in cells with a highly elevated PomY-mCh concentration. Stunningly, not only PomX/PomY-mCh clusters but also the “pure” PomY-mCh clusters were proficient in determining the site of cell division (Fig. 4D), demonstrating that cell division site positioning solely depends on the presence of a PomY condensate, irrespectively of the presence of PomX.

PomX interacts with full-length PomY; specifically, PomX^CC^ interacts with PomY, while we have not detected interactions between PomX^IDR^ and PomY (Schumacher et al., 2017; Schumacher et al., 2021). Because PomY at physiological concentrations depends on PomX for condensate formation, we mapped in more detail how PomY interacts with PomX. To this end, we purified the three individual PomY domains as well as combinations of these domains with mCh- and His_6_-tags (from hereon PomY^CC^-mCh, PomY^Heat^-mCh, PomY^IDR^-mCh, PomY^CC-Heat^-mCh and PomY^Heat-IDR^-mCh) as well as a Strep-tagged full-length PomX variant (Fig. S2B; cf. Fig. 2A). In *in vitro* pull-down assays, we observed that PomX-Strep interacted not only with full-length PomY-mCh but also with all the truncated PomY-mCh variants (Fig. 5).

**Fig. 5.**
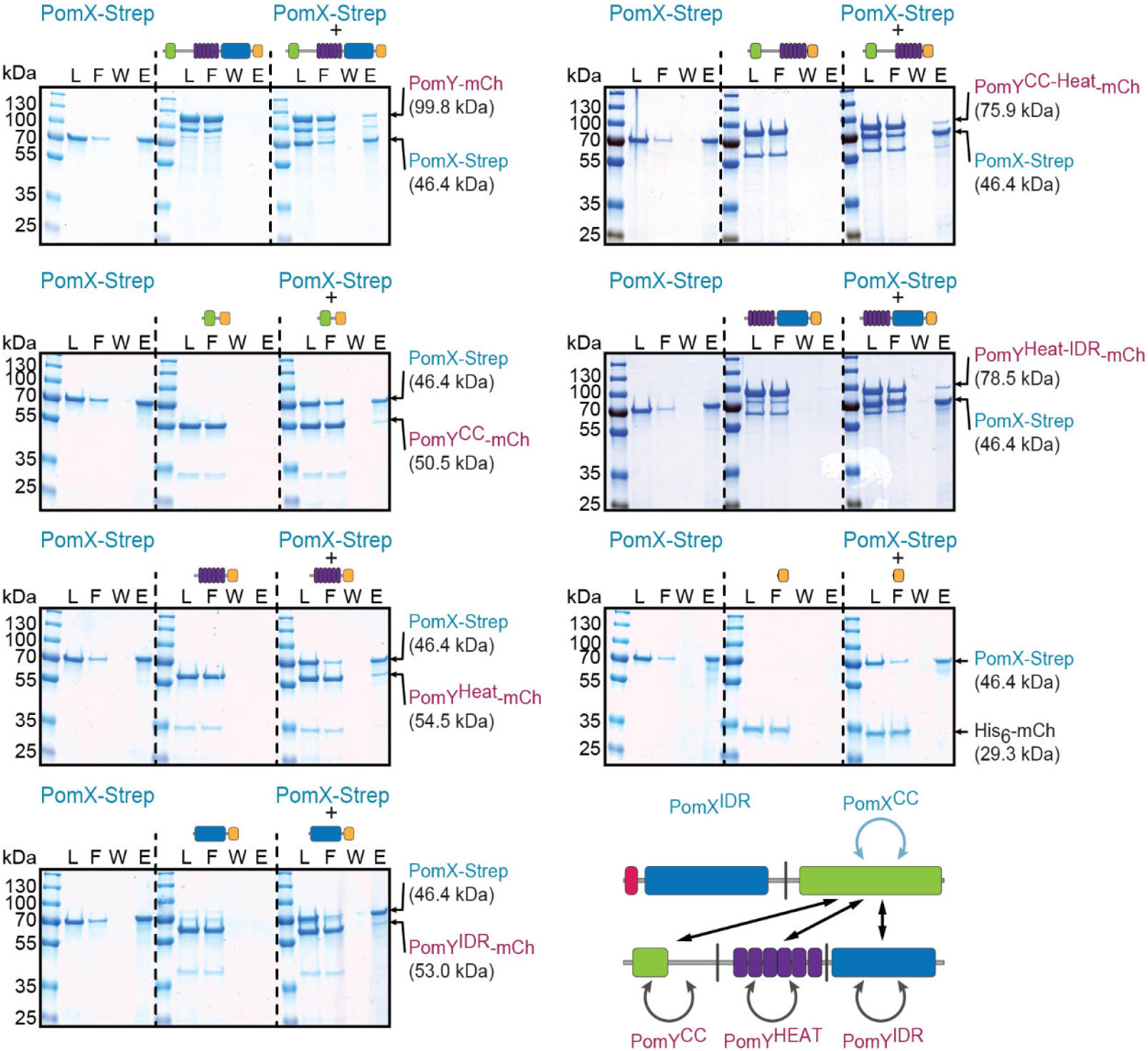
All three PomY domains interact with PomX. *In vitro* pull-down experiments with PomX-Strep and PomY-mCh variants. Proteins were pulled-down using MagStrep XT beads with 10μM proteins alone or in combination as indicated. Proteins from the load (L), flow-through (F), last wash (W), and elution fraction (E) were separated by SDS-PAGE and stained with Instant Blue. All samples in a panel were loaded on the same gel. Stippled lines are included for clarity. Experiments were repeated three times, and representative results are shown. The lower panel summarizes interactions observed in the pull-down experiments (black), the BACTH analyzes of PomY (grey) and in (Schumacher et al., 2021) (blue). See also Figure S2.

Altogether, we conclude that homotypic interactions among PomY molecules are sufficient to drive phase separation with the formation of biomolecular condensates *in vivo*, if the cellular PomY concentration surpasses *C*_sat_. These condensates are functional and support cell division. At physiological concentrations PomY does not form biomolecular condensates independently of PomX. Therefore, we infer that PomX is a scaffold that drives condensate formation by PomY *in vivo* when the cellular PomY concentration is below *C*_sat_ for LLPS. In this mechanism, PomX enriches PomY locally to a concentration above *C*_sat_ via heterotypic interactions involving all three PomY domains. This model is in agreement with PomX filaments nucleating PomY condensate formation via surface-assisted condensation. As cells contain only a single large PomX scaffold, PomY condensate formation is restricted to the single position of the PomX scaffold.

### Multivalent homotypic PomY interactions and multivalent heterotypic interactions with PomX are required for PomY condensate formation *in vivo*

Next, we dissected PomY to understand how the PomY condensates are formed and to identify the domains that mediate or contribute to condensate formation *in vitro* and *in vivo.* To test for condensate formation *in vitro*, we used the *in vitro* LLPS assay with varying concentrations of the PEG8000 crowding agent together with the same mCh- and His6-tagged PomY variants as in the pull-down experiments. We found that all constructs containing PomY^CC^ formed condensates in a PEG8000 dependent manner (Fig. 6AB). Neither PomY^HEAT^, PomY^IDR^ nor both combined formed condensates, but both domains stimulated condensate formation by PomY^CC^ (Fig. 6AB). We conclude that *in vitro* homotypic interactions between PomY^CC^ domains are required and sufficient for PomY condensate formation and that the two additional domains stimulate phase separation.

**Fig. 6.**
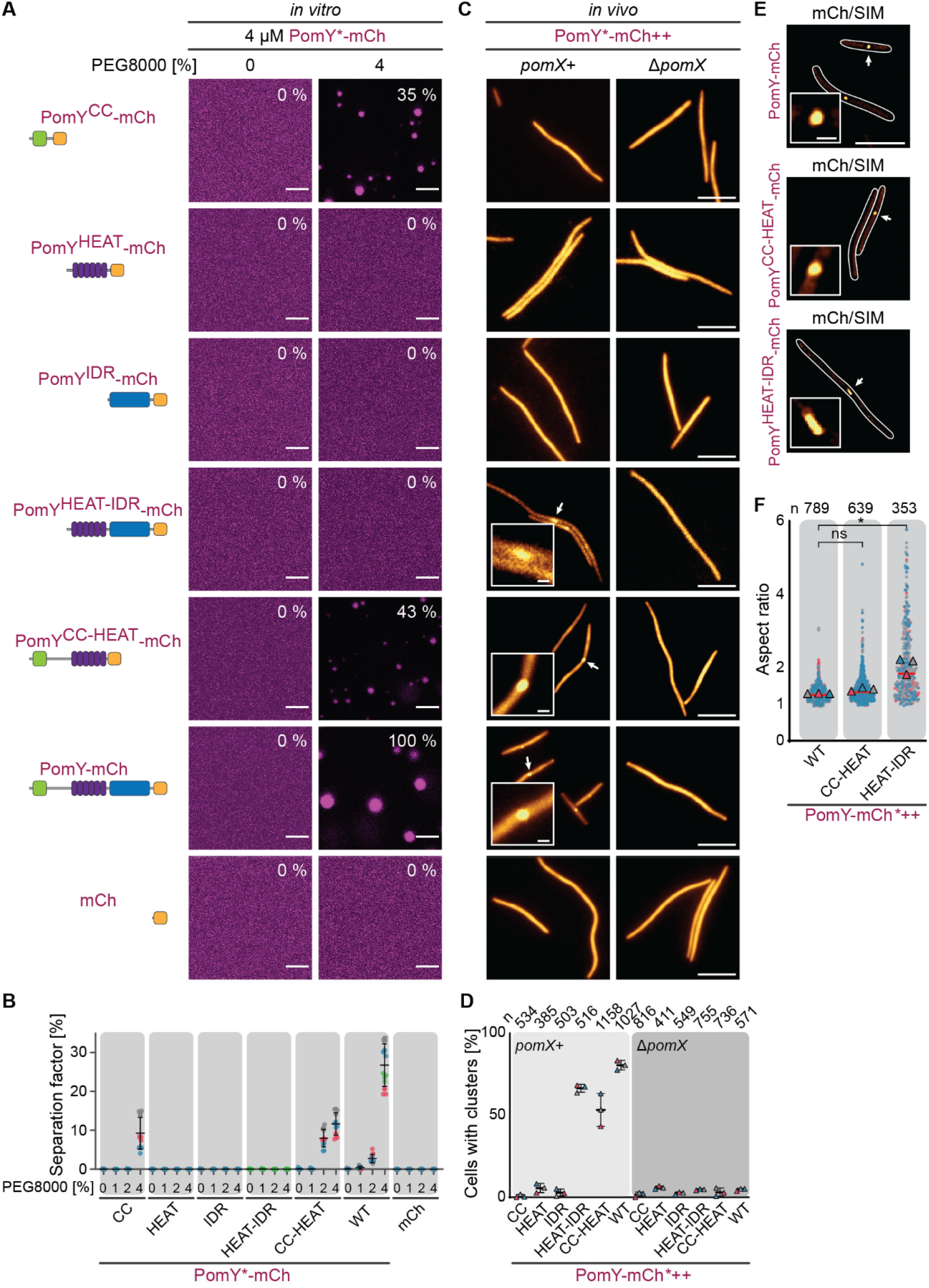
Multivalent homotypic PomY interactions and multivalent heterotypic interactions with PomX are required for PomY condensate formation *in vivo*. A, B. PomY^CC^ is necessary and sufficient for phase separation *in vitro*, while PomY^HEAT^ and PomY^IDR^ stimulate phase separation. A. Representative images of PomY-mCh variants in the presence and absence of 4% PEG8000 *in vitro.* Average separation factor in % normalized to the separation factor of PomY-mCh is indicated in the images and calculated from the maximum intensity projection of confocal z-stacks of the individual conditions. Scale bars, 5μm. B. Analysis of the separation factor of PomY-mCh variants *in vitro* in the presence of different PEG8000 concentrations. The average separation factor in % is calculated from the maximum intensity projection of confocal z-stacks of the individual conditions. Data from at least three independent experiments each shown as colored dots with >10 analyzed images per condition. Error bars indicate mean±STDEV. C. Fluorescence microscopy of cells expressing the PomY-mCh variants indicated in A. White arrows, clusters shown in the insets. ++ indicates that PomY-mCh variants were expressed from the *pilA* promoter. Scale bars, 5μm and 0.5μm in insets. D. % of cells in C with a cluster. Cells from three independent experiments were analyzed and the mean of each experiment shown as a colored triangle, error bars indicate mean±STDEV. E. SIM images of cells expressing PomY-mCh variants as in C.White arrows indicate clusters shown in the insets. Scale bars, 5μm and 0.5μm in inset. F. Aspect ratio of clusters of the indicated PomY-mCh variants based on SIM images from E. Measurements from three independent experiments are shown as colored dots together with the mean. Red line indicates the median. The number of analyzed clusters (n) is indicated on top. *, *P*<0.01, ns, not significant in 2way ANOVA test. See also Figures S2 and S5.

To delineate which PomY domains are important for condensate formation *in vivo*, we expressed the same domains and combinations thereof fused to mCh in the presence and absence of PomX using the strong, constitutively active *pilA* promoter. The truncated proteins accumulated at levels equal to or moderately higher than that of PomY-mCh expressed from its native promoter (with the possible exception of PomY^HEAT^-mCh, which accumulated, but the levels are unknown) (Fig S5A). None of the truncations complemented the *ΔpomY* cell length and division defects, demonstrating that all three domains are required for full PomY function (Fig. S5B). PomY^CC-HEAT^- and PomY^HEAT-IDR^-mCh, almost as efficiently as PomY-mCh, formed a single cluster per cell in a PomX-dependent manner, while cluster formation was largely abolished for the other truncated variants (Fig. 6C). None of the individual domains formed a cluster, demonstrating that the interactions between individual PomY domains and PomX are of low affinity and insufficient to drive cluster formation. These findings are in agreement with cluster formation being mediated by multiple heterotypic interactions between PomX and PomY at native protein levels.

The observation that PomY^CC^ did not form a cluster *in vivo*, even though it undergoes LLPS *in vitro*, albeit to a lower extent than the WT protein, suggests that the expression level *in vivo* is insufficient to surpass *C*_sat_ even in the presence of PomX. Surprisingly, PomY^HEAT-IDR^ formed a cluster *in vivo*, even though this variant lacks PomYCC required for phase separation *in vitro.* High-resolution SIM demonstrated that PomY-mCh and PomY^CC-HEAT^-mCh (both phase separate *in vitro*) clusters were spherical to spheroidal, while PomY^HEAT-IDR_^-mCh (does not phase separate *in vitro*) clusters were strongly elongated (Fig. 6EF, cf. Fig. 1B). The spherical shape of the PomY^CC-HEAT^-mCh clusters supports that these clusters, similar to those of PomY-mCh, reflect surface-assisted condensate formation by PomY on the PomX scaffold. By contrast, the highly elongated shape of the PomY^HEAT-IDR^ clusters suggests that this protein also does not phase separate *in vivo* but instead interacts with the PomX scaffold without undergoing phase separation with condensate formation. We conclude that not the presence but the shape of the cellular PomY clusters indicates phase separation. In summary, our data support that cluster formation by PomY *in vivo* is a two-pronged mechanism: (1) multivalent heterotypic interactions between PomX and PomY domains locally enrich PomY on the PomX scaffold, and (2) homotypic, multivalent PomY interactions enable phase separation, and, thus, surface-assisted condensate formation on the PomX scaffold below Csat; for (2) to occur, the homotypic interactions between PomY^CC^ domains are essential. We also speculate that the surface tension of the PomY condensates formed on the PomX scaffold is likely to drive their reorganization into the spheroidal condensates observed *in vivo.*

### PomY condensates enrich FtsZ and nucleate GTP-dependent FtsZ polymerization

FtsZ is recruited by the PomX/PomY complex to form the Z-ring and initiate cell division, while FtsZ in *ΔpomY* cells localizes in a speckled pattern and cell division occurs away from the PomX cluster (Schumacher et al., 2017). Having found that the “pure” cellular PomY condensates in the absence of PomX are sufficient for cell division site positioning, we sought to determine how they accomplish this function.

To gain insights into how PomY condensates stimulate Z-ring formation, we asked whether PomY condensates can recruit FtsZ using a Cys-tagged FtsZ (Cys-FtsZ) labeled with Alexa488 (A488-FtsZ) in our *in vitro* system. In control experiments, we found that purified Cys-FtsZ behaved as untagged FtsZ and sedimented in a GTP-dependent manner in high-speed centrifugation experiments and had GTPase activity (Fig. S6ABC) suggesting that it polymerizes in a GTP-dependent manner to form filaments. Next, we formed PomY-mCh condensates in the presence of the PEG8000 crowding agent and A488-FtsZ or a commericial Streptavidin Alexa488 conjugate (Strep-A488). A488-FtsZ partitioned into the PomY-mCh condensates and was significantly more enriched than the Strep-A488 negative control (Fig. 7AB), while phase separation by PomY-mCh was not influenced by the presence of A488-FtsZ or Strep-A488 (Fig. S6D). Importantly, neither A488-FtsZ nor Strep-A488 formed visible structures or condensates in the absence of PomY-mCh (Fig. 7A). Thus, PomY condensates concentrate FtsZ *in vitro.*

**Fig. 7.**
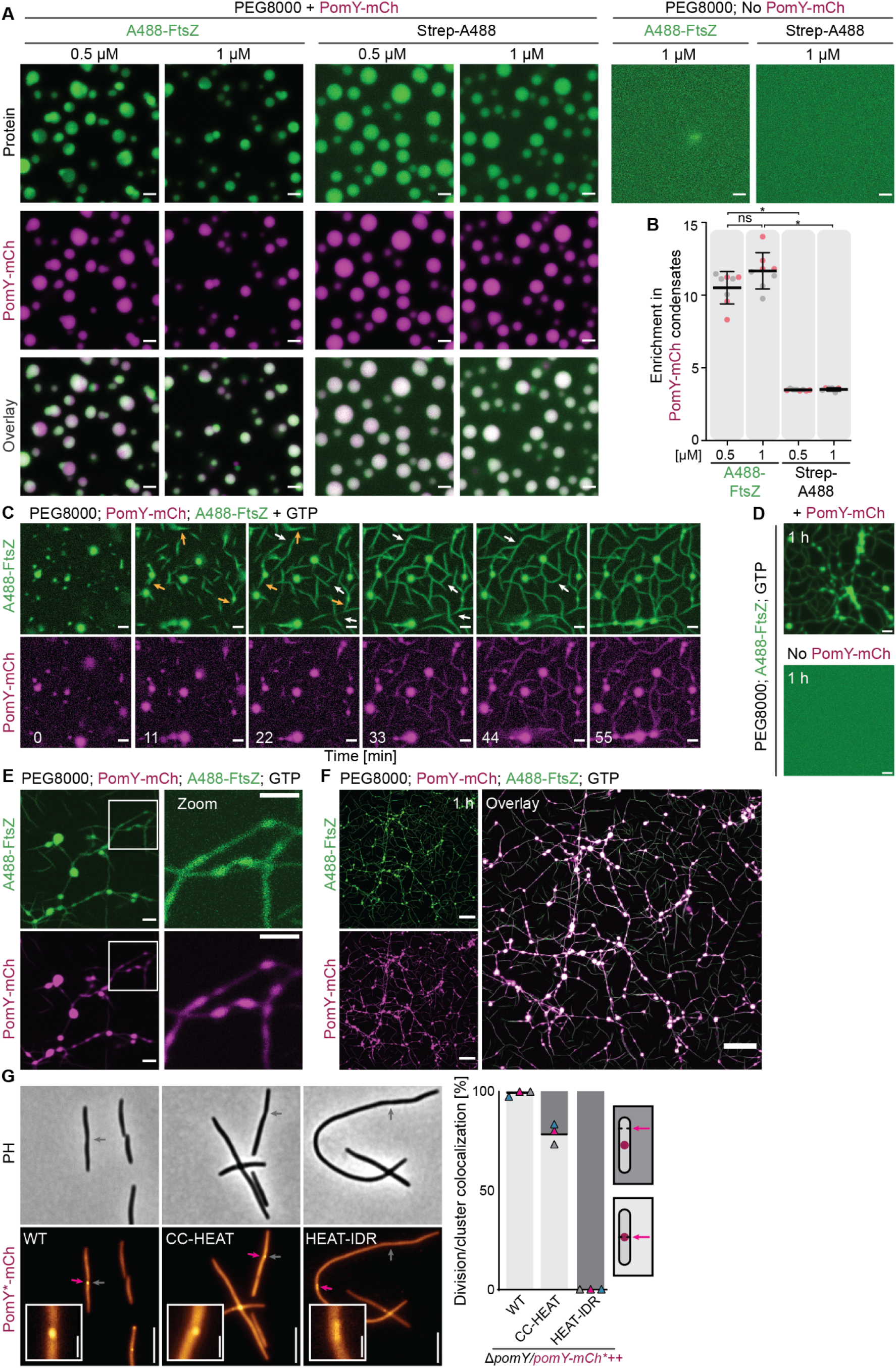
PomY condensates enrich FtsZ and nucleate GTP-dependent FtsZ polymerization. A, B. FtsZ selectively partitions into PomY-mCh condensates, but is unable to form condensates by itself in presence of crowding agent. A. Representative images of either 1 μM A488-FtsZ or Strep-A488 in presence of 4% PEG8000 and in presence or absence of 4 μM PomY-mCh. Scale bars, 5μm. B. Quantification of the enrichment of A488-FtsZ and Strep-A488 in PomY-mCh condensates based on the experiments in A. Magenta and grey points are from two independent experiments with n=8 analyzed images per condition. Error bars indicate mean ± STDEV. *, *P*<0.0001, ns, not significant in 2way ANOVA multiple comparisons test. C-F. FtsZ polymerizes from PomY-mCh condensates, bundle and align and deform PomY-mCh condensates. C. Time-series of 1μM A488-FtsZ in the presence of 4μM PomY-mCh, 2% PEG8000 and 0.5 mM GTP. Images were acquired close to the coverslip surface. Orange arrows indicate A488-FtsZ filament bundles emerging from condensates. White arrows indicate fusion and alignment of A488-FtsZ filament bundles. The images have been gamma-corrected for better visualization. Unaltered images are included in Fig. S6E. Scale bars, 5μm. D. Maximum intensity projection of a confocal z-stack of 1μM A488-FtsZ in presence of 2% PEG8000 in the presence or absence of 4μM PomY-mCh after 1h of incubation. Scale bar, 5μm. E. High-magnification images of PomY-mCh and A488-FtsZ under conditions as in C. Scale bars, 5μm. F. Maximum intensity projection of a confocal z-stack of A488-FtsZ in the presence or absence of PomY-mCh after 1h of incubation under the conditions in in C and D. Scale bar, 25μm. G. Quantification of the colocalization of clusters of PomY-mCh variants with division constrictions. Representative images of cells with cell division constrictions (grey arrows) and PomY*-mCh clusters (magenta arrow). Histogram, black lines indicate the mean of three independent experiments shown as triangles in different colors. Light grey and dark grey, indicate constrictions colocalizing with a PomY*-mCh cluster or not colocalizing, respectively. ++ indicates that proteins were expressed from the *pilA* promoter. Scale bars, 5μm and 1μm in insets. See also Figure S6 and Movie 3.

To assess whether PomY condensates could act as nucleation centers for FtsZ polymerization, we induced PomY-mCh condensate formation at 2% PEG8000 in the presence of A488-FtsZ and GTP. Remarkably, filamentous FtsZ structures emerged from PomY-mCh condensates within minutes, later on forming an extended filamentous network (Fig. 7C, Fig. S6E; Movie 3). These A488-FtsZ structures are μm-wide and extend up to several μm in length, suggesting that they represent bundles of individual FtsZ filaments, which are only one monomer wide, i.e. 5-7nm, and 50-100nm long and cannot be resolved by fluorescence microscopy (Hou et al., 2012; Mukherjee and Lutkenhaus, 1998). Indeed under these conditions, but in the absence of PomY-mCh, A488-FtsZ did not generate visible structures at this optical resolution (Fig. 7D). PomY-mCh condensates were often deformed into lenticular shapes in this process and appeared to cover the FtsZ filamentous structures (Fig. 7E) and after more than an hour of incubation, PomY-mCh colocalizes with the whole FtsZ network underscoring the liquid nature of PomY condensates (Fig. 7F). These larger networks are formed by fusion and alignment of A488-FtsZ filamentous structures emerging from distinct PomY-mCh condensates (Fig. 7C, Fig. S6E). We conclude that PomY condensates specifically enrich FtsZ, and serve as nucleation centers for GTP-dependent FtsZ polymerization and bundling of FtsZ filaments into larger structures.

To investigate the link between PomY phase separation, FtsZ recruitment, and cell division *in vivo*, we took advantage of PomY variants that form a cluster in the presence of PomX and can (PomY-mCh, PomY^CC-HEAT^-mCh) or cannot undergo phase separation (PomY^HEAT-IDR^-mCh) (Fig. 6C-F). We analyzed these variants for their ability to recruit FtsZ and stimulate Z-ring formation by scoring the colocalization of PomY*-mCh clusters and cell division constrictions (Fig. 7G). Similar to PomY-mCh, PomY^CC-HEAT^-mCh clusters mostly colocalized with division constrictions, suggesting that this variant can recruit FtsZ and stimulate Z-ring formation. By contrast, PomY^HEAT-IDR^-mCh clusters did not colocalize with constriction sites. Although we cannot rule out that PomY^CC^, in addition to being central for condensate formation *in vitro* and *in vivo*, is also central for the interaction to FtsZ, these observations suggest that PomY phase separation is required to recruit FtsZ to the PomY/PomX/PomZ complex to stimulate Z-ring formation. We also note that phase separation by PomY is not sufficient to stimulate cell division to WT levels as PomY^CC-HEAT^ does not complement the *ΔpomY* mutant (Fig. S5BC). In total, our data show that PomY condensates act as a concentration and nucleation hub for FtsZ, and strongly support that this is the underlying mechanism by which Z-ring formation is stimulated precisely over the PomX/PomY/PomZ cluster.

## Discussion

Here, we identified a new mode of cell division regulation in bacteria via formation of a selective biomolecular condensate. This condensate emerges from the PomY protein undergoing LLPS based on multivalent interactions with itself and the molecular PomX scaffold. The PomX scaffold locally enriches PomY and thus allows the formation of the PomY condensate below *C*_sat_ for LLPS. Thereby this PomX-mediated surface-assisted condensation restricts condensate formation to a single position in the cell. This localized PomY condensate, in turn, selectively enriches the tubulin homolog FtsZ, the key protein in bacterial cell division, to nucleate and bundle GTP-dependent FtsZ filaments, thereby guiding Z-ring formation and subsequent cell division precisely over the PomX/PomY cluster.

That PomY undergoes LLPS to form a biomolecular condensate is supported by several observations, all hallmarks of LLPS (Alberti and Hyman, 2021; Banani et al., 2017; Lyon et al., 2021; Shin and Brangwynne, 2017). *In vivo* and at physiological concentrations, the PomX-dependent PomY cluster is spherical to spheroidal, non-stoichiometric, scales in size with cell size, exchanges molecules with the cytoplasm, disintegrates during division and is subsequently formed *de novo*. Finally, at very high cellular concentrations PomY by itself forms spherical clusters. *In vitro* PomY forms spherical, liquid-like condensates in a protein and crowding agent concentration-dependent manner. Moreover, phase separation both *in vitro* and *in vivo* is driven by multivalent homotypic interactions. Specifically, PomY^CC^ is both necessary and sufficient for LLPS *in vitro*, while PomY^HEAT^ and PomY^IDR^ stimulate LLPS. Similar domains have previously been implicated in phase separation behaviour (Borcherds et al., 2021; Woodruff et al., 2017; Yoshimura and Hirano, 2016). The many charged residues and the unusually high isoelectric point of PomY together with the strong salt dependence of PomY LLPS and its sensitivity to 1,6-hexanediol *in vitro* all suggest that electrostatic interactions between charged sequence motifs, as well as hydrophobic interactions contribute to its phase separation.

In contrast to PomY, and despite the overall similar structural features of PomX and PomY, PomX does not phase separate. Rather we confirmed that PomX self-assembles into filaments that bundle at high concentrations *in vitro* (Schumacher et al., 2017; Schumacher et al., 2021) and that associate into a highly stable, filamentous scaffold *in vivo.* This structure undergoes fission during cell division and then doubles in size over the cell cycle by incorporating PomX molecules.

Why then do two distinct mechanisms of macromolecular complex formation exist within the PomX/PomY cluster, filament formation and phase separation, and how do their interplay lead to the formation of a single complex *in vivo?* In absence of PomX, PomY forms condensates only when the bulk concentration is above *C*_sat_. *In vitro C*_sat_ is surpassed either by elevation of protein concentration or addition of crowding agents that by an excluded volume effect are believed to increase the effective protein concentration (Kuznetsova et al., 2014) (Fig. 8AB). Analogously, *in vivo*, PomY only forms condensates independent of PomX if it is highly overexpressed, suggesting that native PomY levels, where PomY by itself is homogenously distrubuted, are well below *C*_sat_. Importantly, in presence of PomX, PomY also forms structures below *C*_sat_. *In vitro*, PomY was enriched on the PomX filaments, coated and bundled these structures below *C*_sat_; above *C*_sat_, PomY condensates also formed in the bulk and wetted the PomX filaments. Similarly, *in vivo* PomY only forms a single cluster per cell which colocalizes with the PomX scaffold (Schumacher et al., 2017) at native concentrations, *i.e.* below *C*_sat_; when PomY is overexpressed it can also form multiple ones, suggesting that some of them form independently of PomX (Fig. 8B).

**Fig. 8.**
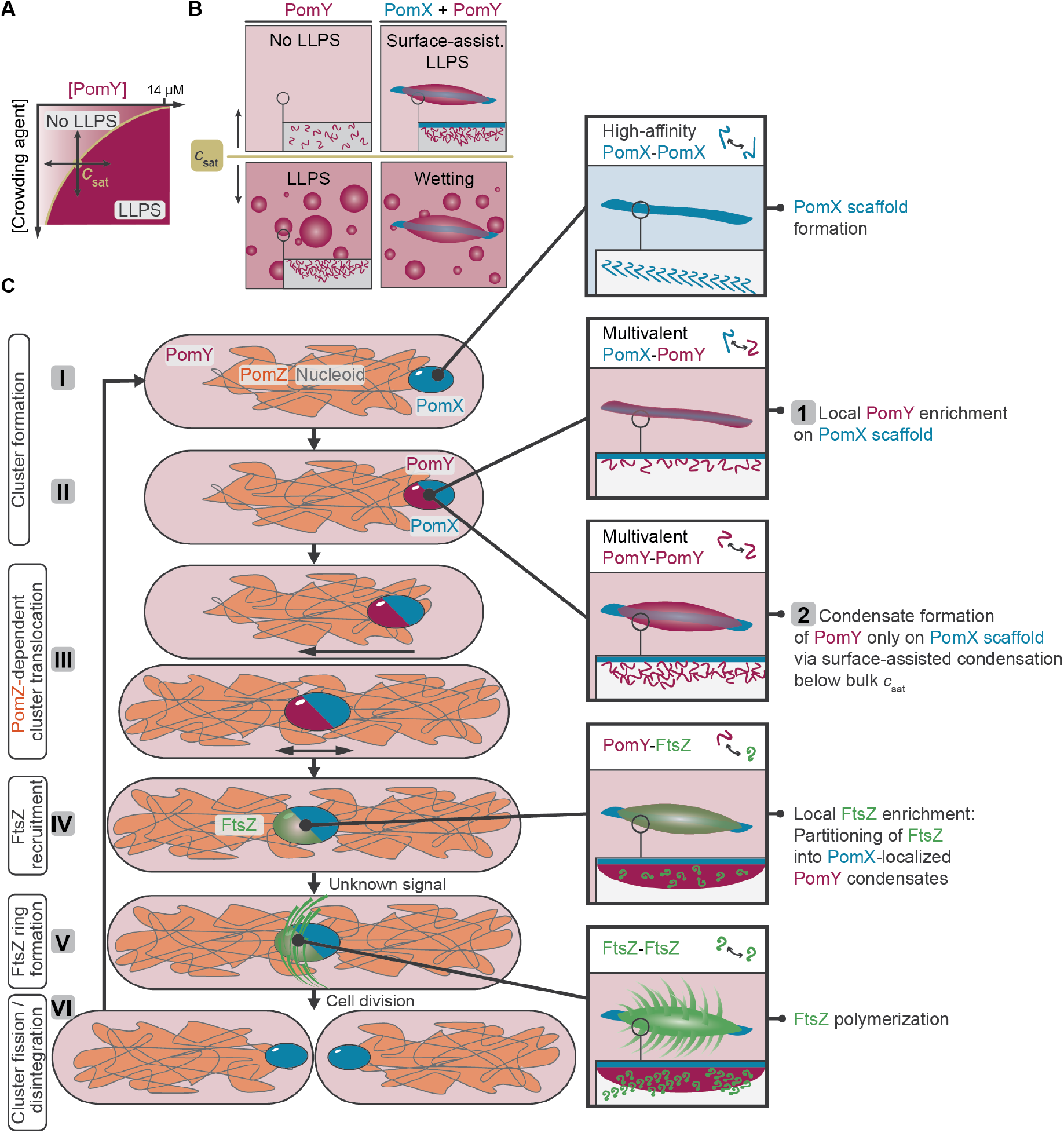
A phase-separated biomolecular PomY condensate nucleates polymerization of the tubulin homolog FtsZ to spatiotemporally regulate bacterial cell division. See main text for details

Altogether, this implies that the PomX/PomY cluster at physiological concentrations is formed by surface-assisted condensation of PomY on the PomX scaffold, which proceeds in two steps as suggested by dissection of PomY. First, PomX interacts with all three PomY domains, and the opposing isoelectric points of PomX (pI 4.9) and PomY (pI 9.3) suggest electrostatic interactions between the proteins. These multivalent heterotypic interactions between PomX and PomY locally enrich PomY on the PomX scaffold resulting in a high local PomY concentration. Second, because the PomX-dependent PomY clusters display all the hallmarks of biomolecular condensates, we infer that multivalent, homotypic interactions between PomY domains enable phase separation of PomY locally on the PomX scaffold. Importantly, because the bulk concentration is well below *C*_sat_ for LLPS, PomY forms a single condensate precisely on the PomX scaffold. Similar surface-assisted condensate formation has been observed in the spatiotemporal regulation of eukaryotic protein condensates on microtubules, membranes and DNA (Gouveia et al., 2022; King and Petry, 2020; Morin et al., 2022; Rouches et al., 2021; Setru et al., 2021; Siahaan et al., 2019; Snead et al., 2022; Tan et al., 2019; Zhao et al., 2021). Specifically, it has been suggested that surface-assisted condensation can occur via prewetting (Morin et al., 2022; Rouches et al., 2021; Zhao et al., 2021). Our data indicate that prewetting also underlies the formation of the PomX/PomY complex observed here. Of note, we observe that PomY is more spheroidal than PomX in the PomX/PomY cluster, supporting that PomY indeed forms a single condensate on the PomX scaffold below *C*_sat_, that reorganizes into spherical shapes due to its surface tension. However, *in vitro* we observed PomY coating the PomX filaments in a seemingly uniform layer below *C*_sat_. This discrepancy can be explained by the extended length of the PomX structures *in vitro*, which are very different from those observed *in vivo.* Moreover, we did not observe this PomY layer breaking up into more round condensed phases akin to the (regular) droplet formation of condensates coating fibers or filaments based on hydrodynamic instability (King and Petry, 2020; Morin et al., 2022; Rayleigh, 1878; Setru et al., 2021). FRAP recovery of the PomY condensates on a minute time scale suggests that they have a high viscosity which might only lead to minor variations in PomY density that cannot be resolved by our optical resolution as the underlying hydrodynamic instability depends on viscosity and surface tension (Morin et al., 2022; Setru et al., 2021).

PomY condensates *in vitro* enrich the tubulin homolog FtsZ and nucleate GTP-dependent FtsZ polymerization and bundling of FtsZ filaments. The nucleation of FtsZ filament bundles by PomY condensates is phenomenonologically similar to synthetic coacervates that artificially enrich FtsZ (te Brinke et al., 2018). More importantly, it is strikingly reminiscent of eukaryotic condensates that stimulate polymerization of cytoskeletal proteins like tubulin (Hernandez-Vega et al., 2017; King and Petry, 2020; Woodruff et al., 2017) or actin (Banjade and Rosen, 2014; Graham et al., 2022; Li et al., 2012; Su et al., 2016; Yang et al., 2022), suggesting a broadly conserved role of condensates in the spatiotemporal regulation of protein filament formation and, in particular, that condensate stimulation of tubulin polymerization is an ancient mechanism. *In vitro* PomY condensates deformed during FtsZ filament formation and covered the FtsZ filament bundles leading to the alignment and bundling of growing filaments into larger networks. However, whether bundling of FtsZ filaments by PomY has a functional role *in vivo* remains to be uncovered. Nevertheless, we speculate that PomY could act as a transient crosslinker that bundles FtsZ similar to *E. coli* ZapA (Caldas et al., 2019).

Taken together, we propose a novel model for regulating bacterial cell division by a biomolecular condensate (Fig. 8C). High-affinity interactions among PomX molecules drive assembly of a single filamentous PomX scaffold per cell (Fig. 8C, I). During the cell cycle, additional PomX molecules are incorporated into this scaffold allowing it to double in size. PomY associates with the PomX scaffold via multivalent heterotypic interactions, thereby locally reaching a high concentration resulting in condensate formation on the PomX scaffold based on homotypic PomY interactions (Fig. 8C, II). The PomY condensate exchanges molecules with the cytoplasm, but overall there is a net gain of PomY molecules in the condensate allowing it to grow over time. The PomX/PomY complex translocates on the nucleoid towards midcell via PomX/PomY-stimulated ATP hydrolysis by the DNA-bound PomZ ATPase (Schumacher et al., 2017; Schumacher et al., 2021) (Fig. 8C, III). At midcell, the PomY condensate recruits FtsZ by direct interaction and enriches FtsZ locally (Fig. 8C, IV). Upon an unknown signal, FtsZ filaments emerge from the PomY condensates to form the Z-ring consisting of treadmilling FtsZ filaments on the membrane (Fig. 8C, V).

Subsequently, FtsZ recruits other proteins required to execute cytokinesis. During cytokinesis, the PomX scaffold undergoes fission with both daughters “receiving” a smaller scaffold, while the PomY condensate disintegrates (Fig. 8C, VI). We speculate that the advantage of the PomX scaffold fission is that the daughters are each born with a single PomX scaffold, thereby ensuring that only a single PomY condensate forms. Similarly, we suggest that PomY condensate disintegration represents an elegant mechanism to prevent the immediate stimulation of additional divisions after a completed division event. Altogether, this mechanism represents a *bona fide* positive mechanism for spatiotemporal regulation of Z-ring formation and cell division in bacteria. We speculate that a similar mechanism might be employed by analogous positive regulatory systems, especially in unusually shaped bacteria since it does not require cellular symmetry as in negative regulatory systems (Kiekebusch et al., 2012; Ramm et al., 2019).

Our study raises several intriguing questions for future research avenues. First, we do not know how the remarkable fission of the PomX scaffold occurs during cell division. Our data support that this event does not involve a duplication event, unlike centrioles that duplicate before segregation (Pelletier et al., 2004). PomX scaffold splitting follows a cell division-specific signal (Schumacher et al., 2017; Schumacher et al., 2021), which we suspect is of mechanical nature, e.g. septum ingrowth. Second, the mechanism underlying PomY condensate disintegration also remains unknown. Even though disintegration occurs less often at increased PomY concentration (Schumacher et al., 2017), it is unlikely to be caused by dilution of PomY because total cell volumes do not change during division, as opposed to the sudden volume increase during nuclear envelope breakdown in eukaryotes that results in disintegration of the nucleolus condensates (Lafontaine et al., 2021). We speculate that PomY condensate disintegration could be triggered by other cell division proteins disrupting PomX-PomY or PomY-PomY interactions. A sudden increase of the surface-to-volume ratio of the PomY condensate as a consequence of the PomX scaffold fission could also lead to PomY condensate disintegration. Alternatively, the association of PomY with FtsZ filaments as we observed *in vitro* could influence PomY condensate stability. Third, we cannot explain why the PomY condensate only becomes competent for stimulating Z-ring formation at midcell. Maturation of PomY condensates, or the PomZ ATPase might allow Z-ring stimulation only at midcell. Alternatively, the nucleoid, similar to nucleoid occlusion effects in other systems, could influence Z-ring formation. Finally, it remains to be uncovered how the PomZ ATPase associates with the PomX/PomX complex to stimulate its translocation on the nucleoid to midcell.

Interestingly, AAPs of other ParA/MinD-type ATPases, *i.e.* ParB and McdB, have recently been shown to undergo phase separation (Babl et al., 2022; Guilhas et al., 2020; MacCready and Vecchiarelli, 2021). Although PomY, ParB and McdB share little sequence homology, these observations suggest that surface-assisted condensate formation by AAPs of MinD/ParA-like ATPase is a general mechanism of these proteins in orchestrating the spatiotemporal organization of bacteria.

## Acknowledgements

We thank Giovanni Cardone (MPIB Imaging Facility) and Silke Bergeler for assistance with image analysis, Leopold Urich (MPIB Core Facility) for assistance with the operation of the liquid handler, Gabriele Malengo (MPITM Core Facility for Flow Cytometry and Imaging) for help with structured illumination microscopy, Kerstin Andersson and Katharina Nakel for assistance with protein purification, and Lei Kai, Philipp Blumhardt and Leon Babl for helpful discussions. B.R. was supported by a fellowship from the German Research Council (DFG) through the Graduate School of Quantitative Biosciences Munich and in part by the National Science Foundation grant no. PHY-1734030. P.S. and L.S.-A. acknowledge financial support by the DFG through the Transregio 174 “Spatiotemporal dynamics of bacterial cells’’ (Project ID No. 269423233) and the Max Planck Society. P.S. acknowledges the support of the research network MaxSynBio via a joint funding initiative of the German Federal Ministry of Education and Research (BMBF).

## Author contributions

B.R. and D.S. contributed equally to this work. The order of the two shared first authors is alphabetical and has no further meaning. B.R., D.S., P.S. and L.S.-A. conceived the study. B.R. and D.S designed *in vitro* experiments, D.S. and P.K. designed *in vivo* experiments. B.R., D.S., T.H., A.H., P.K., and F.M. performed experiments. B.R. and D.S. analyzed data and wrote the original manuscript draft. B.R., D.S., P.S. and L.S.-A. reviewed and edited the manuscript. All authors discussed the results and revised the manuscript.

## Competing interests

The authors declare no competing interests.

## Methods

Detailed Methods are provided in Supplementary Information.

## Notes

### Competing Interest Statement

The authors have declared no competing interest.

