## Supplementary Information for "A phase-separated biomolecular condensate nucleates polymerization of the tubulin homolog FtsZ to spatiotemporally regulate bacterial cell division"

**This file contains**

- Supplementary Figures S1-S6
- Supplementary Tables S1-S3
- Legends to Movies 1-3
- Methods
- Supplementary References

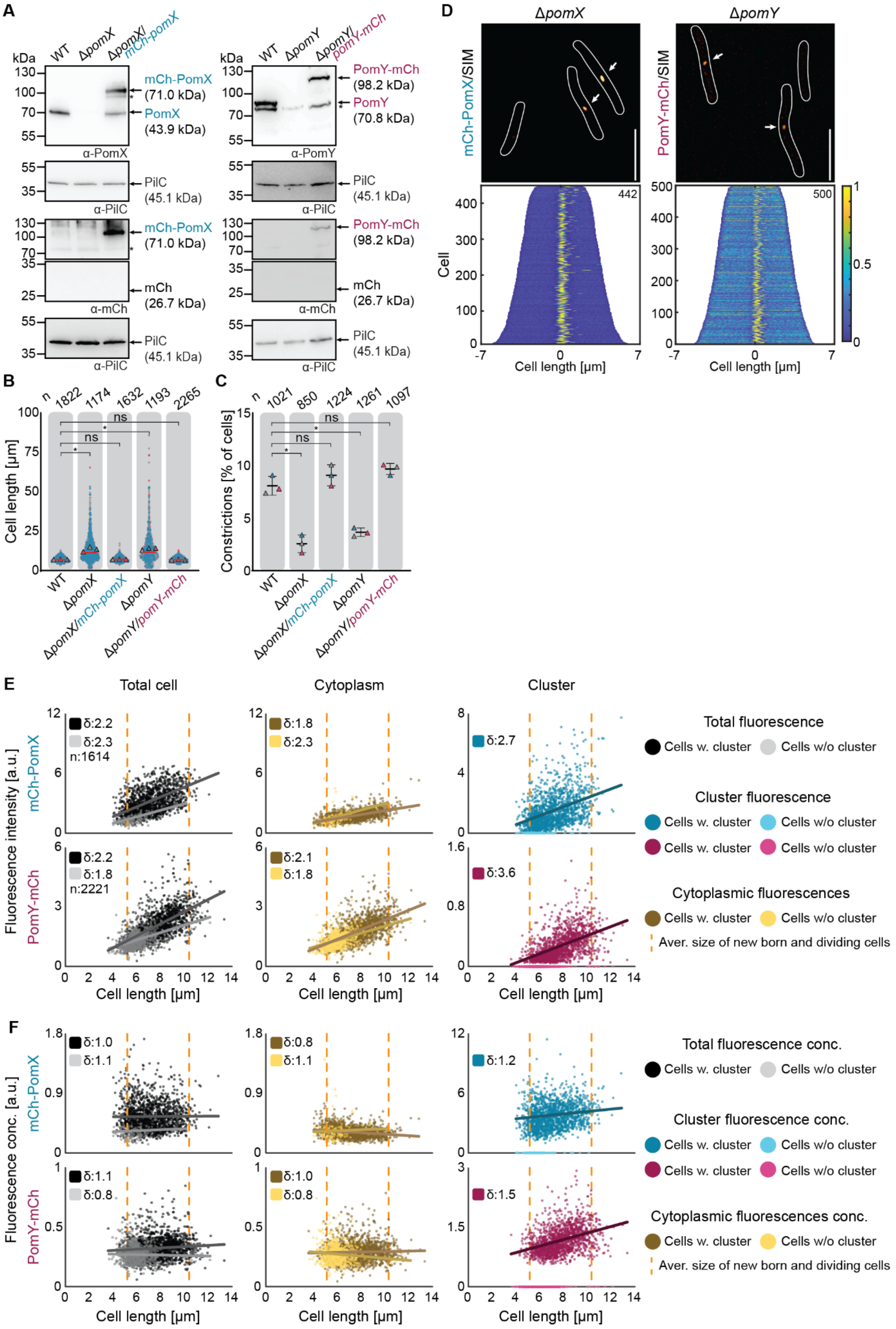
A
kDa
WT
ΔpomX
ΔpomX/mCh-pomX
130
100
70
55
mCh-PomX
(71.0 kDa)
PomX
(43.9 kDa)
α-PomX
PilC
(45.1 kDa)
α-PilC
mCh
(26.7 kDa)
α-mCh
ΔpomY
ΔpomY/pomY-mCh
PomY-mCh
(98.2 kDa)
PomY
(70.8 kDa)
α-PomY
D
mCh-PomX/SIM
PomY-mCh/SIM
Cell
Cell length [µm]
442
500
B
n
1822
1174
1632
1193
2265
Cell length [µm]
ns
WT
ΔpomX
ΔpomX/mCh-pomX
ΔpomY
ΔpomY/pomY-mCh
C
1021
850
1224
1261
1097
Constrictions [% of cells]
E
Total cell
Cytoplasm
Cluster
Fluorescence intensity [a.u.]
mCh-PomX
PomY-mCh
δ:2.2
δ:2.3
n:1614
δ:1.8
δ:2.3
δ:2.7
δ:2.2
δ:1.8
n:2221
δ:2.1
δ:1.8
δ:3.6
Cell length [µm]
Total fluorescence
Cells w. cluster
Cells w/o cluster
Cluster fluorescence
Cytoplasmic fluorescences
Aver. size of new born and dividing cells
F
Fluorescence conc. [a.u.]
δ:1.0
δ:1.1
δ:0.8
δ:1.1
δ:1.2
δ:1.1
δ:0.8
δ:1.0
δ:0.8
δ:1.5
Total fluorescence conc.
Cluster fluorescence conc.
Cytoplasmic fluorescences conc.

**Supplementary Figure 1. Native level expression of mCh-PomX and PomY-mCh restores a WT phenotype**

A. Immunoblot analysis of mCh-PomX and PomY-mCh accumulation. The same amount of protein was loaded per lane, separated by SDS-PAGE, blotted, and probed with α-PomX, α-PomY, α-mCh and α-PilC (loading control) antibodies. Molecular size markers are indicated on the left. * indicates unspecific binding of the antibodies used. Proteins with their predicted MW are indicated on the right. Experiments were repeated three times with similar results, and representative data are shown.

B. Cell length distribution of indicated strains. Data from three independent replicates are shown as magenta, grey, and teal. The mean for each experiment is shown as triangles in the relevant color. Red line indicates the median. The number of analyzed cells (n) is shown at the top. *, $P<0.01$, ns, not significant, 2way ANOVA with multiple comparisons to WT.

C. Constrictions frequency analysis of strains in B. Data from three independent replicates are shown as magenta, grey, and teal with the mean±STDEV. *, $P<0.01$, ns, not significant in 2way ANOVA with multiple comparisons to WT.

D. Analysis of cells expressing mCh-PomX and PomY-mCh at native levels by SIM microscopy. White arrows point to mCh-PomX and PomY-mCh clusters. White lines indicate cell outlines. Scale bars, 5µm. Demographs show fluorescence signals of cells sorted according to length and with off-center signals to the right. Number of cells (n) is indicated in the upper right. The colored scale bar indicates fluorescence signal intensity with the highest signal set to 1.

E. Total cellular, cluster, and cytoplasmic fluorescence intensity of cells expressing mCh-PomX (upper panels) and PomY-mCh (lower panels) at native levels as a function of cell length. Orange stippled lines indicate length of average newborn and dividing cell. Dark-colored lines are linear regression. δ values indicate the fold increase between birth and division. Data from three replicates are shown.

F. Total cellular, cluster, and cytoplasmic fluorescence concentrations of mCh-PomX (upper panels) and PomY-mCh (lower panels) in cells of E as a function of cell length. Graphs are as in E.

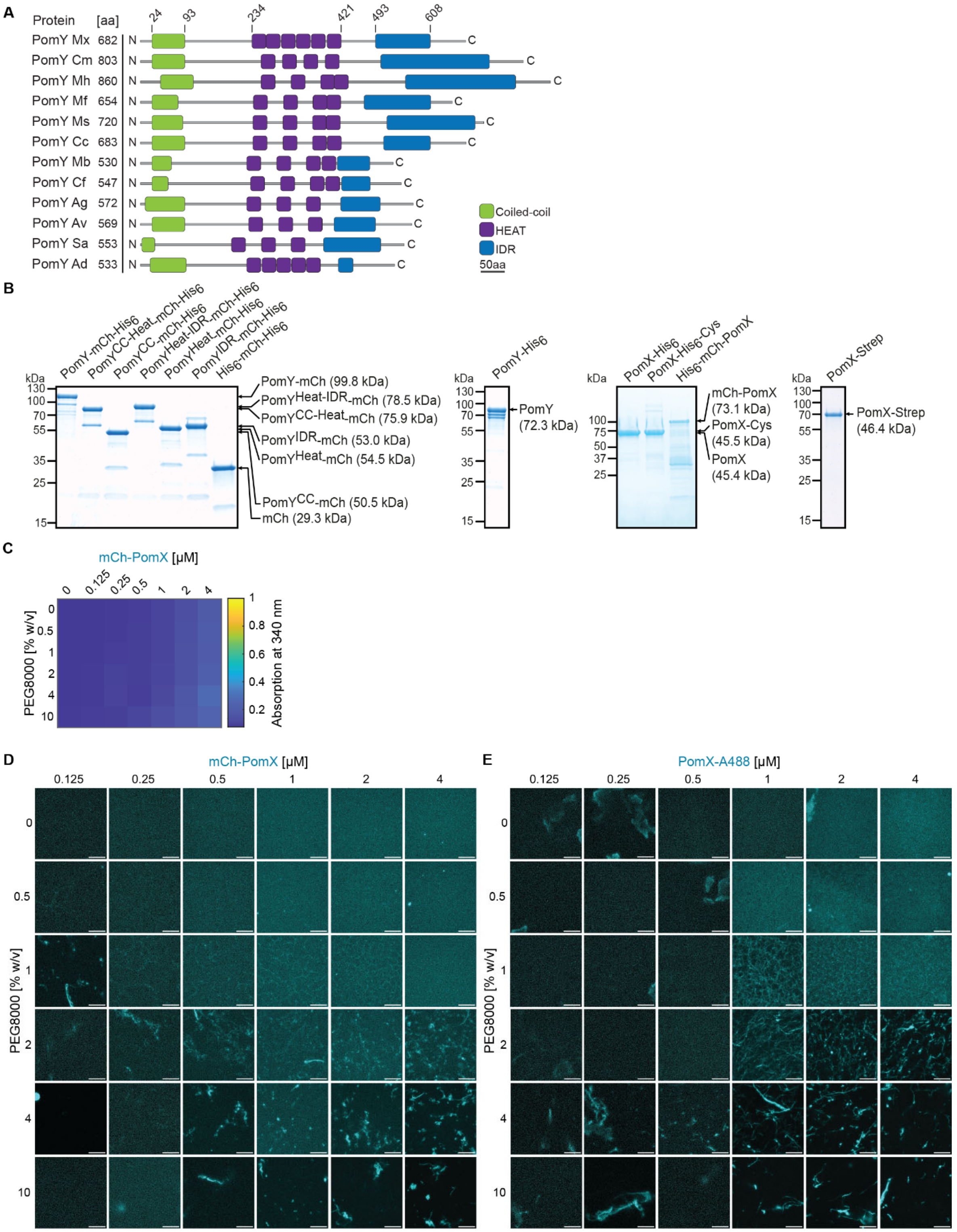


**Supplementary Figure 2. mCh-PomX and PomX form filaments *in vitro*.**
A. Domain analysis of PomY orthologs. PomY orthologs were identified in a best-best hit reciprocal BlastP analysis from fully-sequenced genomes of myxobacteria. Domains prediction was performed using Interpro, disorder prediction using IUPred2A. Numbers on top indicate domain borders.

B. Instant Blue stained SDS-PAGE analysis of purified PomY-mCh-$His_6$ variants, $His_6$-mCh, PomY-$His_6$ and PomX variants. Note that the $His_6$-tagged PomX and PomY variants are referred to as indicated on the right. Molecular size markers are shown on the left. 2µg protein per lane were loaded. Note that PomY-mCh-$His_6$ does not separate to run at the calculated MW in SDS-PAGE, but instead runs at MW of ~120 kDa. Moreover, PomX-$His_6$, PomX-$His_6$-Cys, $His_6$-mCh-PomX and PomX-Strep also do not separate to run at the expected MW in SDS-PAGE, but instead run at MW of ~73 kDa, ~73 kDa, ~100 kDa and 70 kDa, respectively.
C. Turbidity measurements of mCh-PomX for increasing protein and PEG8000 concentrations. Heat map displays average absorption at 340 nm for n=3 replicates.
D, E. PomX forms filaments that are increasingly bundled with increasing concentrations of PEG8000. Representative maximum intensity projections of confocal z-stacks of mCh-PomX (D) and PomX-A488 (E) at different protein and PEG8000 concentrations. Scale bars, 50µm. The images' brightness and contrast levels are not comparable but were individually adjusted for visualization purposes.

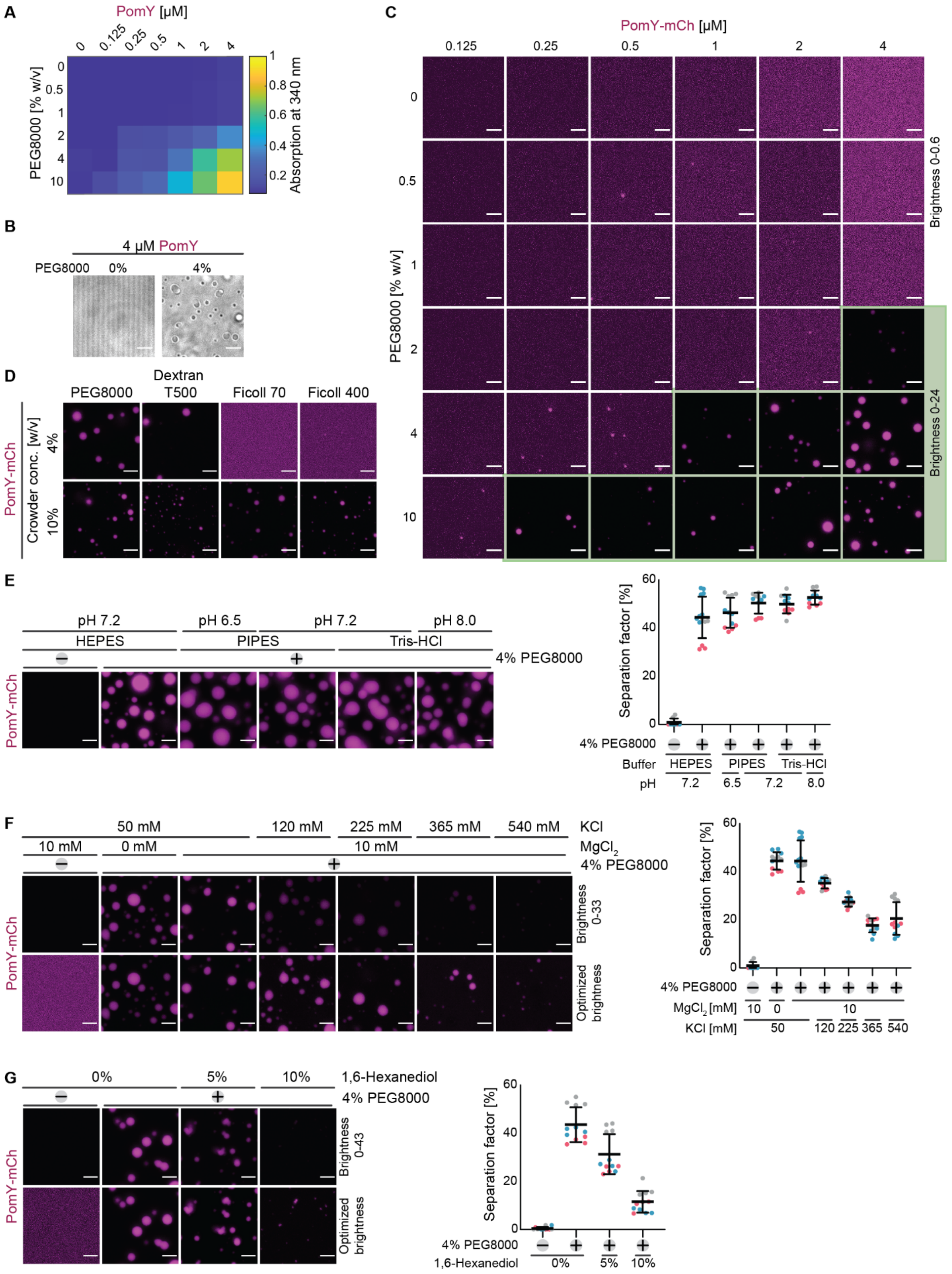


**Supplementary Figure 3. PomY and PomY-mCh phase separate *in vitro*.**
A. Turbidity measurements of PomY for increasing protein and PEG8000 concentrations. The heat map displays average absorption at 340 nm for n=3 replicates.

B. Representative bright-field images of PomY in the absence and presence of PEG8000. Scale bars, 5µm.
C. Representative images of PomY-mCh at different protein and PEG8000 concentrations. Brightness and contrast of the images were adjusted with two different thresholds for clarity indicated by a green background. Scale bars, 5µm.
D. Representative images of 4µM PomY-mCh in the presence of different artificial crowding agents. Scale bars, 5µm.
E. Representative images of PomY-mCh in buffer with different pH based on different buffer substances (4µM PomY-mCh, 0 or 4% PEG8000). The average separation factor (right panel) in percent is calculated from the maximum intensity projection of confocal z-stacks. Right diagram, data shown in magenta, grey and teal originates from three independent experiments with in total >8 analyzed images per condition. Error bars indicate mean ± STDEV.
F. Representative images of PomY-mCh in the absence or presence of 10 mM $MgCl_2$ (left) and in the presence of different KCl concentrations (4µM PomY-mCh, 0 or 4% PEG8000). Top row: brightness and contrast comparable between images, set to 0-33; bottom row: brightness and contrast optimized for each image. The average separation factor in percent is calculated as in (E). Right diagram, data shown in magenta, grey and teal originates from three independent experiments with in total >8 analyzed images per condition. Error bars indicate mean ± STDEV.
G. Representative images of PomY-mCh in the presence of an increasing amount of 1,6-hexanediol (4µM PomY-mCh, 0 or 4% PEG8000). Top row: brightness and contrast comparable between images, set to 0-43; bottom row: brightness and contrast optimized for each image. The average separation factor in percent is calculated as in (E). Right panel, data shown in magenta, grey and teal originates from three independent experiments with in total 12 analyzed images per condition. Error bars indicate mean ± STDEV.

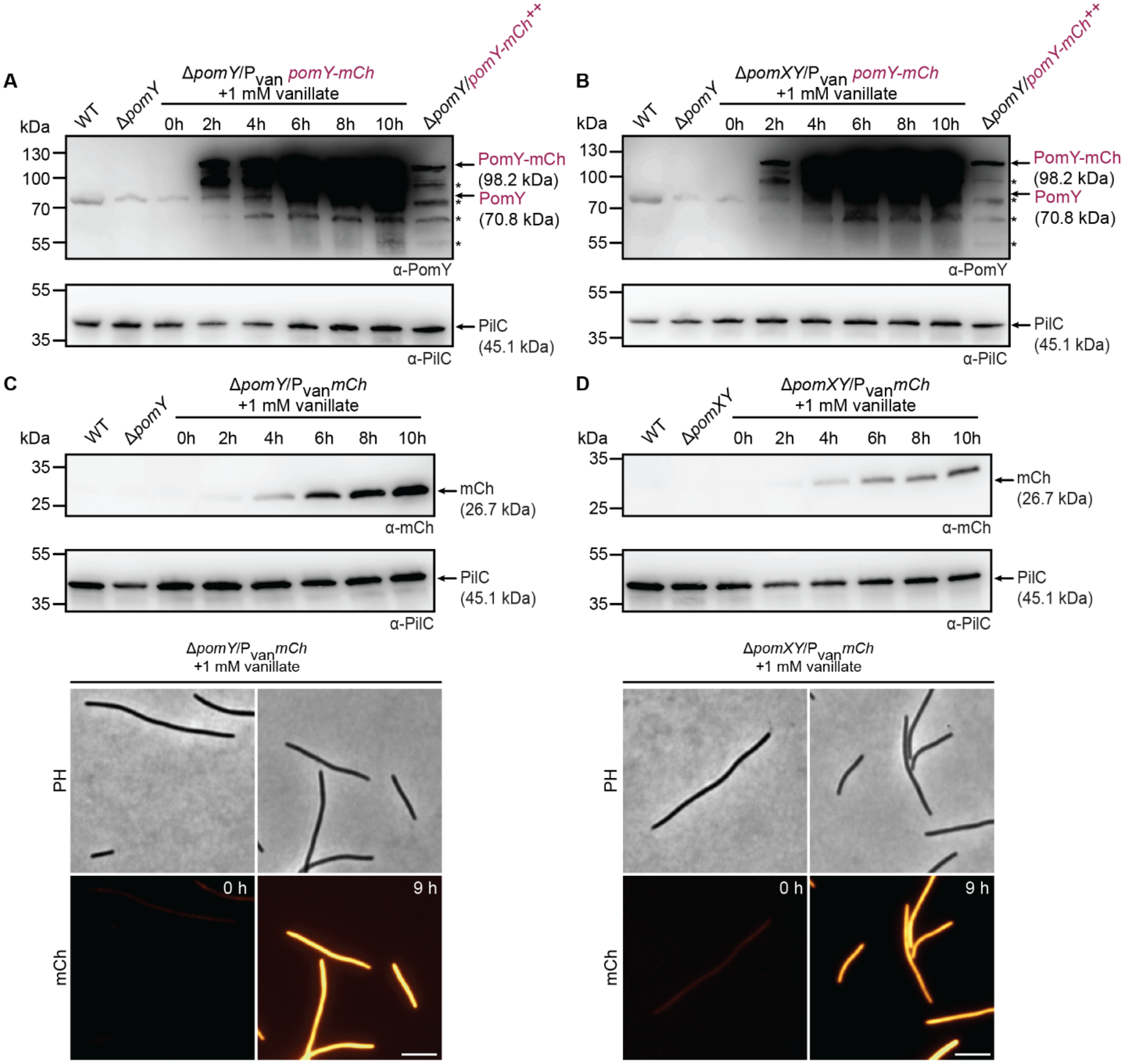


**Supplementary Figure 4. Analysis of strains extensively overproducing PomY-mCh.** A, B. Immunoblot analysis of PomY-mCh accumulation in total cell lysates of indicated strains. The same amount of protein was loaded per lane, separated by SDS-PAGE, blotted, and probed with specific α-PomY and α-PilC (loading control) antibodies. Molecular size markers are indicated on the left. Proteins with their calculated MW are indicated on the right. * indicates unspecific binding of the antibodies used. ++ indicates that these genes were expressed from the constitutively active *pilA* promoter and are included for comparison. Overproduction of PomY-mCh was induced by the addition of 1mM vanillate to cells in suspension at 0h. The experiment was repeated three times, and representative results are shown. Note that PomY-mCh does not separate to run at the expected MW in SDS-PAGE, but instead run at a MW of ~120 kDa.

C, D. Immunoblot analysis of mCh in lysates of indicated strains (upper panel). Blots were done as in A but specific α-mCh antibodies were used. Lower panels, fluorescence microscopy of representative cells at the 0h and 9h time point (lower panel). Scale bar, 5μm.

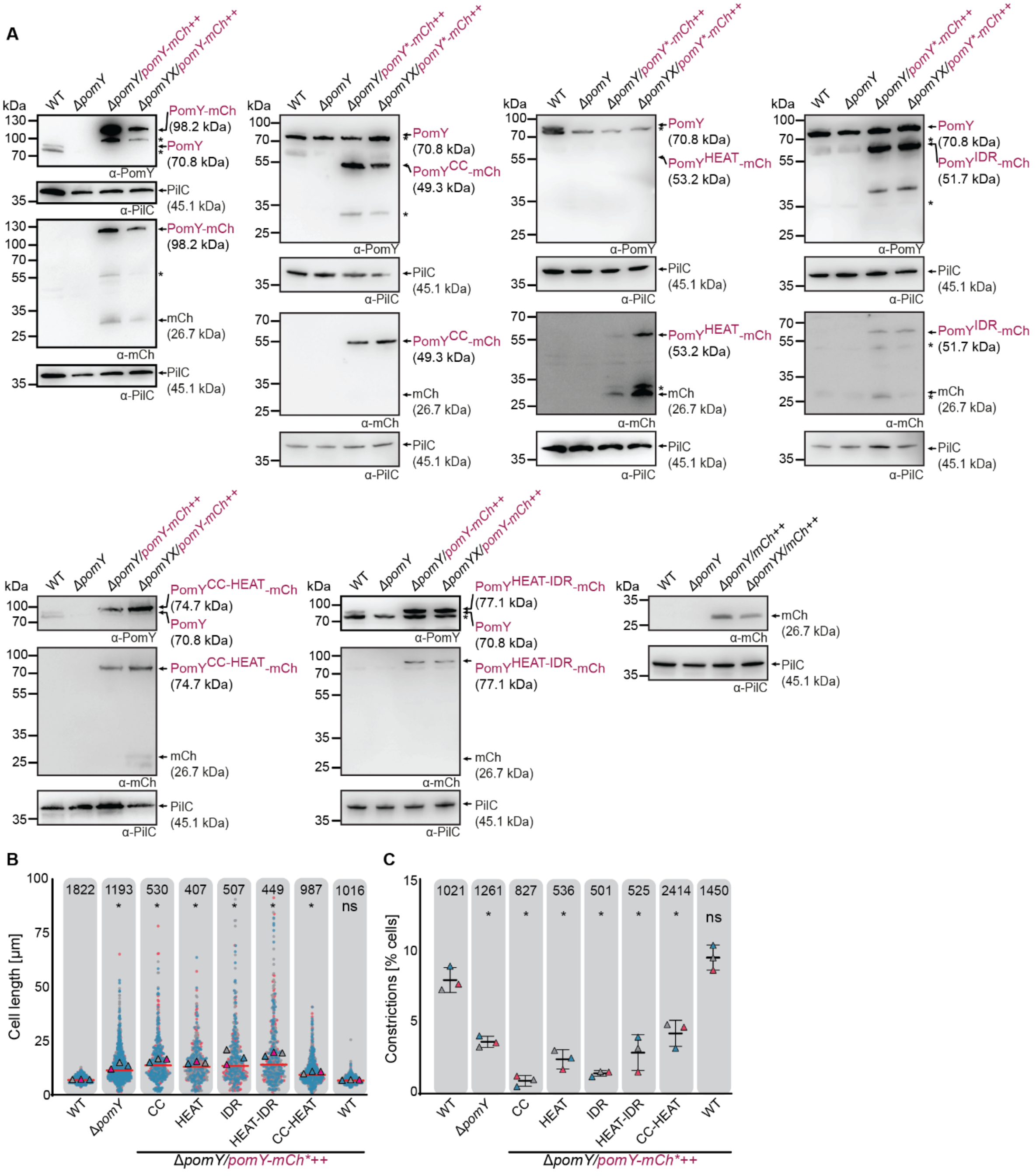


**Supplementary Figure 5. Analysis of truncated PomY-mCh variants**

A. Immunoblot analysis for the accumulation of the indicated PomY-mCh variants, and mCh in strains of the indicated genotype. The same amount of protein was loaded per lane, separated by SDS-PAGE, blotted, and probed with specific α-PomY, α-mCh antibodies and α-PilC (loading control) antibodies. Molecular size markers are indicated on the left. Proteins with their calculated MW are shown on the right. "*" indicates unspecific binding of the antibodies used. The experiments were repeated three times with similar results, and representative data are shown. ++ indicates that these genes were expressed from the *pilA* promoter. Note, that PomY-mCh and PomY$^{IDR}$-mCh do not separate to run at the expected MW in SDS-PAGE, but instead run at MW of ~120 kDa and 60kDa, respectively.

B. Cell length distribution of strains expressing indicated PomY-mCh variants. ++ as in A. Data from three independent replicates are shown as magenta, grey, and teal. The means

are shown as a triangle in the relevant color. Red line shows the median. n are lised at the top. *, *P*<0.01, ns, not significant in 2way ANOVA with multiple comparisons to WT.
C. Constriction frequency analysis of cells of indicated genotypes. ++ as in A.Magenta, grey, and teal triangles show constrictions frequency in % of cells of three independent replicates. Black lines indicate the mean ± STDEV. n are lised at the top. *, *P*<0.01, ns, not significant in 2way ANOVA with multiple comparisons to WT.

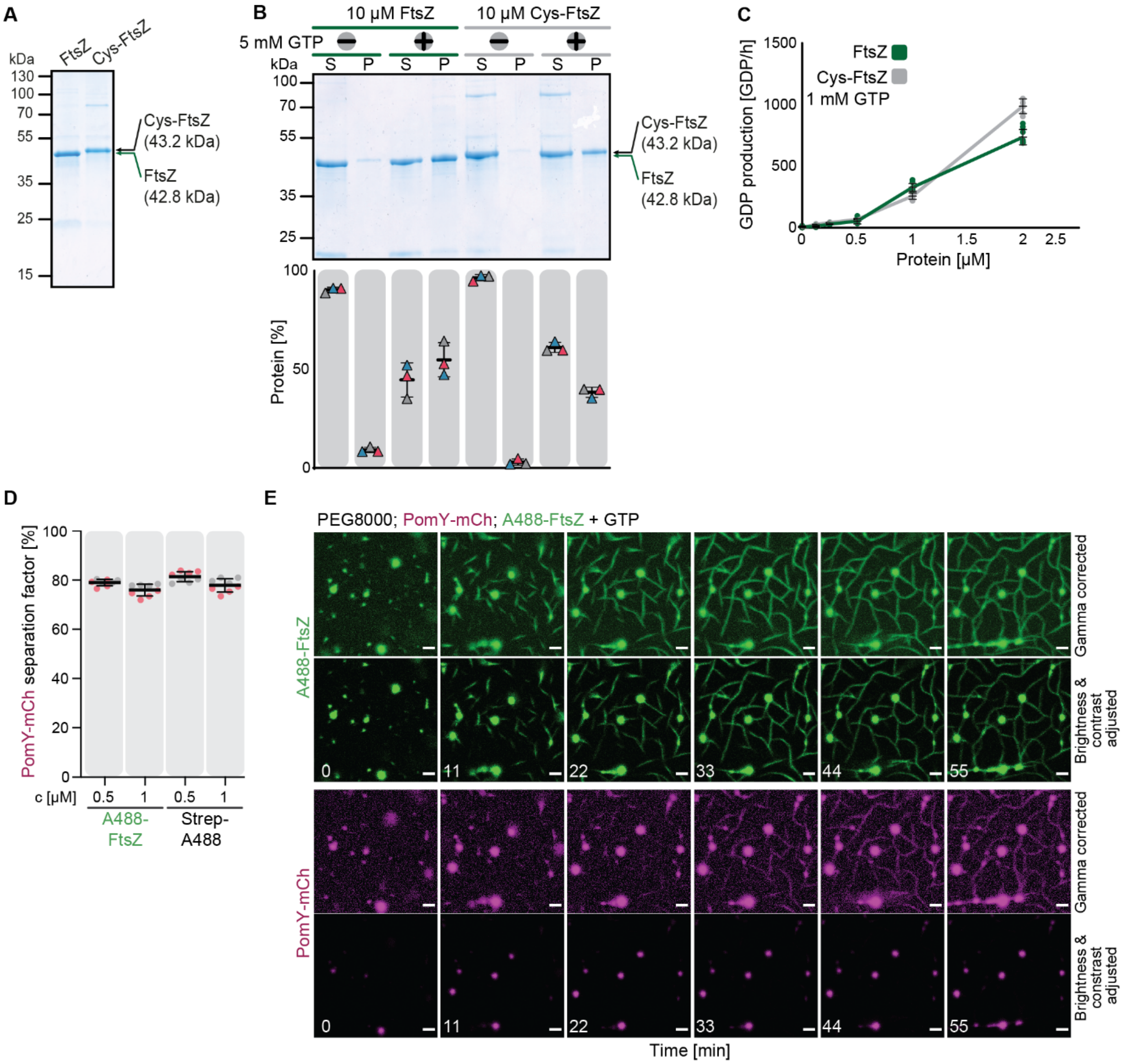


**Supplementary Figure 6. Characterization of Cys-FtsZ and A488-FtsZ**

A. Instant Blue-stained SDS-PAGE analysis of purified FtsZ and Cys-FtsZ. Molecular size markers are shown on the left and proteins with their molecular weight are shown on the right. 2µg protein per lane were loaded

B. Instant Blue-stained SDS-PAGE of soluble and pellet fraction of 10µM Cys-FtsZ (unlabeled) and FtsZ after application to high-speed centrifugation with and without the addition of 5mM GTP (upper panel). Protein fractions of three independent replicates were quantified in % of total protein and plotted as colored triangles as the mean ± STDEV (lower panel).

C. Indicated concentrations of Cys-FtsZ (unlabeled) and FtsZ were assayed for their ability to hydrolyze GTP in the presence of 1mM GTP. Green and grey dots indicate independent replicates. The mean ± STDEV are shown as black lines. n for FtsZ, 6; n for Cys-FtsZ, 9.

D. The presence of A488-FtsZ or Strep-A488 does not alter the extent of PomY phase separation. Quantification of the separation factor of PomY-mCh in the presence of 4% PEG8000 and different concentrations of A488-FtsZ or Strep-A488. Data from two independent experiments with in total n=8 analyzed images per condition after 30min of incubation.

E. Gamma-corrected and just brightness and contrast adjusted time-series of A488-FtsZ filament nucleation from PomY-mCh condensates as shown in Fig. 7C. Images were acquired close to the coverslip surface. Scale bars, 5µm.

**Legends to Movies**

**Movie 1: PomY-mCh condensates undergo fusion and relax into spherical shapes.** Two representative time-series of PomY-mCh condensates undergoing fusion (conditions: 4µM PomY-mCh, 4% PEG8000 w/v). White insets represent the location of the snapshots shown Fig. 2H. Scale bars, 5µm.

**Movie 2: PomY-mCh condensates wet PomX-A488 filaments and deform upon contact.** Time-series of PomY-mCh and PomX-A488 in the presence of 4% PEG8000 as shown in the snapshots in Fig. 3B. Scale bars, 5µm.

**Movie 3: In presence of GTP, A488-FtsZ filament bundles emerge from PomY-mCh condensates and align and fuse to give rise to filamentous networks.**
Time-series of A488-FtsZ in the presence of PomY-mCh and GTP. Images were acquired close to the coverslip surface. White inset represents the location of the snapshots shown Fig. 7C and Fig. S6E. The images have been gamma-corrected for better visualization. Scale bars, 5 µm.

Table S1. *M.xanthus* strains used in this study

| Strain | Genotype | Reference |
|---|---|---|
| SA4420 | Δ*mglA* | (Miertzschke et al., 2011) |
| SA4777 | Δ*mglA*, Δ*pomX* | This study |
| SA4779 | Δ*mglA*, Δ*pomY* | This study |
| SA4797 | Δ*mglA*, Δ*pomX*/*attB*::$P_{pomZ}$ *mCh-pomX* (pAH53) | (Schumacher et al., 2017) |
| SA7000 | Δ*mglA*, Δ*pomY*/*attB*::$P_{pilA}$ *pomY-mCh* (pDS7) | (Schumacher et al., 2017) |
| SA7045 | Δ*mglA*, Δ*pomX*, Δ*pomY* | This study |
| SA7064 | Δ*mglA*, Δ*pomY*/*attB*::$P_{nat}$ *pomY-mCh* (pDS8) | This study |
| SA9734 | Δ*mglA*; Δ*pomX*/*attB*::$P_{pilA}$ *mCh-pomX* (pAH35) | This study |
| SA9764 | Δ*mglA*, Δ*pomY*/*mxan18-19*::$P_{van}$ *pomY-mCh* (pDS331) | This study |
| SA9768 | Δ*mglA*, Δ*pomY*/*attB*::$P_{pilA}$ *pomY*$^{CC}$-*mCh* (pPK29) | This study |
| SA9769 | Δ*mglA*, Δ*pomX*, Δ*pomY*/*attB*::$P_{pilA}$ *pomY-mCh* (pDS7) | This study |
| SA9770 | Δ*mglA*, Δ*pomY*/*attB*::$P_{pilA}$ *pomY*$^{CC-HEAT}$-*mCh* (pPK36) | This study |
| SA9772 | Δ*mglA*, Δ*pomX*, Δ*pomY*/*attB*::$P_{pilA}$ *pomY*$^{CC}$-*mCh* (pPK29) | This study |
| SA9779 | Δ*mglA*, Δ*pomX*, Δ*pomY*/*attB*::$P_{pilA}$ *pomY*$^{CC-HEAT}$-*mCh* (pPK36) | This study |
| SA9784 | Δ*mglA*, Δ*pomX*, Δ*pomY*/*attB*::$P_{nat}$ *pomY-mCh* (pDS8) | This study |
| SA9785 | Δ*mglA*, Δ*pomX*, Δ*pomY*/*mxan18-19*::$P_{van}$ *pomY-mCh* (pDS331) | This study |
| SA9794 | Δ*mglA*, Δ*pomY*/*attB*::$P_{pilA}$ *mCh* (pDS79) | This study |
| SA9795 | Δ*mglA*, Δ*pomX*, Δ*pomY*/*attB*::$P_{pilA}$ *mCh* (pDS79) | This study |
| SA9796 | Δ*mglA*, Δ*pomY*/*mxan18-19*::$P_{van}$ *mCh* (pAH225) | This study |
| SA9797 | Δ*mglA*, Δ*pomX*, Δ*pomY*/*mxan18-19*::$P_{van}$ *mCh* (pAH225) | This study |
| SA11802 | Δ*mglA*, Δ*pomY*/*attB*::$P_{pilA}$ *pomY*$^{HEAT}$-*mCh* (pPK45) | This study |
| SA11803 | Δ*mglA*, Δ*pomX*, Δ*pomY*/*attB*::$P_{pilA}$ *pomY*$^{HEAT}$-*mCh* (pPK45) | This study |
| SA11804 | Δ*mglA*, Δ*pomY*/*attB*::$P_{pilA}$ *pomY*$^{IDR}$-*mCh* (pPK46) | This study |
| SA11805 | Δ*mglA*, Δ*pomX*, Δ*pomY*/*attB*::$P_{pilA}$ *pomY*$^{IDR}$-*mCh* (pPK46) | This study |
| SA11806 | Δ*mglA*, Δ*pomY*/*attB*::$P_{pilA}$ *pomY*$^{HEAT-IDR}$-*mCh* (pPK47) | This study |
| SA11807 | Δ*mglA*, Δ*pomX*, Δ*pomY*/*attB*::$P_{pilA}$ *pomY*$^{HEAT-IDR}$-*mCh* (pPK47) | This study |

Table S2. Plasmids used in this study

| **Plasmid** | **Genotype** | **Reference** |
|---|---|---|
| pKA3 | Overexpression of *pomZ-His$_6$* in *E. coli*; Amp$^R$ | (Treuner-Lange et al., 2013) |
| pKA28 | P$_{nat}$ *pomZ-mCh*; *attP*; Tc$^R$ | (Treuner-Lange et al., 2013) |
| pKA45 | P$_{pilA}$ *pomZ-mCh*; *attP*; Km$^R$ | (Schumacher et al., 2017) |
| pKA70 | Overexpression of *ftsZ* in *E.coli*; Amp$^R$ | (Treuner-Lange et al., 2013) |
| pDS3 | Overexpression of *pomY-His$_6$* in *E. coli*; Km$^R$ | (Treuner-Lange et al., 2013) |
| pDS7 | P$_{pilA}$ *pomY-mCh*; *attP*; Km$^R$ | (Schumacher et al., 2017) |
| pDS8 | P$_{nat}$ *pomY-mCh*; *attP*; Tet$^R$ | (Schumacher et al., 2017) |
| pDS37 | IPTG-dependent expression of *mCh-pomX* in *E. coli*; Km$^R$ | (Schumacher et al., 2017) |
| pDS43 | IPTG-dependent expression of *pomZ-mCh* in *E. coli*; Km$^R$ | (Schumacher et al., 2017) |
| pDS46 | IPTG-dependent expression of *pomY-yfp* in *E. coli*; Km$^R$ | (Schumacher et al., 2017) |
| pDS79 | P$_{pilA}$ *mCh*; *attP*; Km$^R$ | This study |
| pDS120 | BACTH plasmid containing *pomY* (pUT18C); Amp$^R$ | (Schumacher et al., 2021) |
| pDS121 | BACTH plasmid containing *pomY* (pKT25); Km$^R$ | This study |
| pDS122 | BACTH plasmid containing *pomY* (pUT18); Amp$^R$ | (Schumacher et al., 2021) |
| pDS123 | BACTH plasmid containing *pomY* (pKNT25); Km$^R$ | This study |
| pDS132 | P$_{pilA}$ *pomY$^{CC-HEAT}$-mCh; attP;* Km$^R$ | This study |
| pDS261 | BACTH plasmid containing *pomY$^{CC}$* (pUT18); Amp$^R$ | This study |
| pDS262 | BACTH plasmid containing *pomY$^{CC}$* (pUT18C); Amp$^R$ | This study |
| pDS263 | BACTH plasmid containing *pomY$^{CC}$* (pKNT25); Km$^R$ | This study |
| pDS264 | BACTH plasmid containing *pomY$^{CC}$* (pKT25); Km$^R$ | This study |
| pDS265 | BACTH plasmid containing *pomY$^{HEAT}$* (pUT18); Amp$^R$ | This study |
| pDS266 | BACTH plasmid containing *pomY$^{HEAT}$* (pUT18C); Amp$^R$ | This study |
| pDS267 | BACTH plasmid containing *pomY$^{HEAT}$* (pKNT25); Km$^R$ | This study |
| pDS268 | BACTH plasmid containing *pomY$^{HEAT}$* (pKT25); Km$^R$ | This study |
| pDS269 | BACTH plasmid containing *pomY$^{IDR}$* (pUT18); Amp$^R$ | This study |
| pDS270 | BACTH plasmid containing *pomY$^{IDR}$* (pUT18C); Amp$^R$ | This study |
| pDS271 | BACTH plasmid containing *pomY$^{IDR}$* (pKNT25); Km$^R$ | This study |
| pDS272 | BACTH plasmid containing *pomY$^{IDR}$* (pKT25); Km$^R$ | This study |

| | | |
|---|---|---|
| pDS331 | $P_{van}$ *pomY-mCh*; *mxan18-19*; $Km^R$ | This study |
| pEMR1 | Overexpression of *pomY-His$_6$* in *E. coli*; $Km^R$ | (Schumacher et al., 2021) |
| pEMR3 | Overexpression of *pomX-His$_6$* in *E. coli*; $Km^R$ | (Schumacher et al., 2017) |
| pAH14 | Overexpression of $His_6$-*mCh;* $Amp^R$ | This study |
| pAH35 | $P_{pilA}$ *mCh-pomX; attP;* $Km^R$ | (Schumacher et al., 2017) |
| pAH53 | $P_{mxan0635}$ *mCh-pomX; attP;* $Km^R$ | (Schumacher et al., 2017) |
| pAH180 | Overexpression of Cys-*ftsZ* in *E. coli*; $Km^R$ | This study |
| pAH187 | Overexpression of *pomY$^{CC-HEAT}$–mCh*-$His_6$ in *E. coli*; $Km^R$ | This study |
| pAH194 | Overexpression of *pomY-mCh*-$His_6$ in *E. coli*; $Km^R$ | This study |
| pAH195 | Overexpression of *pomY$^{HEAT-IDR}$-mCh*-$His_6$ in *E. coli*; $Km^R$ | This study |
| pAH196 | Overexpression of *pomY$^{CC}$-mCh*-$His_6$ in *E. coli*; $Km^R$ | This study |
| pAH197 | Overexpression of *pomY$^{HEAT}$–mCh*-$His_6$ in *E. coli*; $Km^R$ | This study |
| pAH198 | Overexpression of *pomY$^{IDR}$-mCh*-$His_6$ in *E. coli*; $Km^R$ | This study |
| pAH205 | Overexpression of *pomX-Strep* in *E. coli*; $Km^R$ | This study |
| pAH225 | $P_{van}$ *mCh; mxan18-19*; $Km^R$ | This study |
| pPK29 | $P_{pilA}$ *pomY$^{CC}$-mCh; attP;* $Km^R$ | This study |
| pPK30 | $P_{pilA}$ *pomY$^{HEAT}$-mCh; attP;* $Km^R$ | This study |
| pPK36 | $P_{pilA}$ *pomY$^{CC-HEAT}$-mCh; attP;* $Km^R$ | This study |
| pPK45 | $P_{pilA}$ *pomY$^{HEAT}$-mCh; attP;* $Km^R$ | This study |
| pPK46 | $P_{pilA}$ *pomY$^{IDR}$-mCh; attP;* $Km^R$ | This study |
| pPK47 | $P_{pilA}$ *pomY$^{HEAT-IDR}$-mCh; attP;* $Km^R$ | This study |
| pSL16 | Construct for in-frame deletion of *mglA* (*mxan1925*) | (Miertzschke et al., 2011) |
| pMR3690 | Plasmid for vanillate inducible gene expression in *M.xanthus; mxan18-19;* $Km^R$ | (Iniesta et al., 2012) |
| pSWU30-*pomY$^{CC-HEAT}$-mCh* | P*pilA* *pomY$^{CC-HEAT}$*-mCh; *attP*; $Tet^R$ | This study |
| pBR35 | Overexpression of $His_6$-*pomZ-mCh* in *E. coli*; $Amp^R$ | This study |
| pBR36 | Overexpression of $His_6$-*mCh-pomX* in *E. coli*; $Km^R$ | This study |

| | | |
|---|---|---|
| pBR38 | Overexpression of *pomY*-*yfp*-$His_6$ in *E. coli*; $Km^R$ | This study |
| pBR39 | Overexpression of *pomX*-$His_6$-Cys in *E. coli*; $Km^R$ | This study |
| pBR72 | Overexpression of *pomY*-*mCh*-$His_6$ in *E. coli*; $Km^R$ | This study |
| pUT18 | BACTH plasmid; $Amp^R$ | (Karimova et al., 1998) |
| pUT18C | BACTH plasmid; $Amp^R$ | (Karimova et al., 1998) |
| pKT25 | BACTH plasmid; $Km^R$ | (Karimova et al., 1998) |
| pKNT25 | BACTH plasmid; $Km^R$ | (Karimova et al., 1998) |
| pKT25-zip | BACTH plasmid containing leucine zipper of GCN4; $Amp^R$ | (Karimova et al., 1998) |
| pUT18C-zip | BACTH plasmid containing leucine zipper of GCN4; $Km^R$ | (Karimova et al., 1998) |

Table S3. Oligonucleotides used in this work

| Oligonucleotide | Sequence 5'-3' |
|---|---|
| mCherry fwd Start XbaI | GCGTCTAGAATGGTGAGCAAGGGCGAGGAG |
| mCherry stop rev HindIII | GGGAAGCTTTTACTTGTACAGCTCGTC |
| MXAN_0634-6 | GCGTCTAGAGTGAGCGACGAGCGTCCG |
| DS1 | CCGGAATTCCGACGAGCAGTTGAGCACCAG |
| DS2 | GCGAGATCTGGCGGAGCCCCGCGCCCGCACAGGC |
| DS14 | GCGCATATGAGCGACGAGCGTCCG |
| DS40 | GCGGAATTCTTACTTGTACAGCTCGTC |
| KA477 | GCGGGATCCCATGGTGAGCAAGGGCGAGGAG |
| KA478 | GCCAAGCTTTCATTACTTGTACAGCTCGTCCAT |
| AH159 | GGAATTCCATATGGGCAAGTGCAAGGGCTCCGGCGAC CAGTTCGATCAGAACAAGC |
| KA500 | GCGGGATCCTTATTACGGCAGTTCCGTCTGGC |
| NdeI PomY fwd | GGAATTCCATATGAGCGACGAGCGTCCGGAC |
| DS274 | GCGAAGCTTCTTGTACAGCTCGTCCATGC |
| DS12 | GCGCATATGGGACGGTTGGGCGTGAGC |
| DS271 | GCGCATATGCTGGAGCAGCCTCGTCCG |
| DS278 | GCGTCATGAAGAAAGCCTTTGAACAGAACG |
| DS276 | GCGAAGCTTACTTCTCGAACTGTGGGTGACTCCAGCGC ACCGTGGCCTGAC |
| mCherry fwd NdeI | GCGCATATGGTGAGCAAGGGCGAGGAGG |
| CC(nat) linker rev BglII | CGCAGATCTGGCGGACGCCTGCAGGAAGCGCTGCG |
| 17 HEAT(nat) XbaI fwd | GCGTCTAGAGTGGGACGGTTGGGCGTGAGCGC |
| 18 HEAT(nat) linker rev BglII | CGCAGATCTGGCGGACCGCGCCCGCACAGGCTTC |
| DS4 | GCGTCTAGAGTGAGCGACGAGCGTCCG |
| 27 HEAT fwd | GCGTCTAGAGGACGGTTGGGCGTGAGCGC |
| FM4 | GCGTCTAGACTGGAGCAGCCTCGTCCG |
| BR68 | GGCCTGCTGGGTGCC |
| BR70 | ATGGACGAGCTGTACAAGTGATGAAAGCTTGCGGCCG |
| BR64 | CTTGTACAGCTCGTCCATGCC |
| BR69 | CCGTCGCCCAGACGAAG |
| BR71 | TCCGCGGGTCTGGAAGTTCTGTTCCAGGGGCCCAGCAA GGGCGAGGAG |
| BR72 | GTGATGGTGATGGTGATGTTTCATGGTATATCTCCTTATT AAAGTTAAACAAA |
| BR62 | CATGGACGAGCTGTACAAGAAGCTTGCGGCCGCAC |
| BR75 | CATATGTATATCTCCTTCTTAAAGTTAAACAAAATTATTTC TAGAG |
| BR77 | GAAGGAGATATACATATGAGCGACGAGCGTCCG |
| BR87 | GTGGTGGTGGTGGTGGTG |
| BR153 | GGTGGTTCTGGTTGCGGTAGCGGCAGCGGTAGCTGAGA TCCGGCTGCTAACAAAGC |
| BR185 | CTCCAGAACCTCCTCCACCAGCGGCGAAGTATTTGTGCC |
| BR184 | GGTGGAGGAGGTTCTGGAGGCGGTGGAAGTGGTGGCGG AGGTAGCATGGTGAGCAAGGGCGAG |
| Mxan0634 fwd BACTH | GCGTCTAGAGGTGAGCGACGAGCGTCCG |
| Mxan_0634-17 pomY-rev | GCGGGTACCCGAGCGGCGAAGTATTTGTG |
| Mxan0634 Coil KpnI BACTH | GCGGGTACCCGCGCCTGCAGGAAGCGCTG |
| Mxan0634 HEAT XbaI BACTH | GCGTCTAGAGATGGGACGGTTGGGCGTGAGC |

| | |
|---|---|
| Mxan0634 HEAT KpnI BACTH | GCGGGATCCCGCCGCGCCCGCACAGGC |
| Mxan0634 Pro XbaI BACTH | GCGTCTAGAGATGCTGGAGCAGCCTCGTCCG |

**Methods**

M. xanthus and E. coli strains and growth. *M. xanthus* strains are listed in Table S1. Plasmids and oligonucleotides used are listed inTable S2 and S3, respectively. All *M. xanthus* strains are derivatives of DK1622 (WT) (Kaiser, 1979). *M. xanthus* strains were cultivated in 1% CTT medium (1% casitone, 10mM Tris-HCl pH 7.6, 1 mM $KPO_4$ pH 7.6, 8mM $MgSO_4$) or on 1% CTT 1.5% agar plates (Hodgkin and Kaiser, 1977). Kanamycin, oxytetracycline and gentamycin were added at concentrations of 50µg/ml, 10µg/ml and 10µg/ml, respectively. Growth was measured as an increase in optical density (OD) at 550 nm. *M. xanthus* cells were transformed by electroporation with a BioRad MicroPulser™ (BioRad) at 0.65Ω. In-frame deletions were generated as described (Shi et al., 2008). All plasmids were verified by sequencing. All strains were verified by PCR. All *M. xanthus* strains used are non-motile to allow time-lapse microscopy for several hours by deletion of *mglA* (Miertzschke et al., 2011; Schumacher and Søgaard-Andersen, 2018). Plasmids for the expression of genes in *M. xanthus* were integrated by site directed integration in the Mx8 *attB* locus (*attB*) or the *mxan18-19* intergenic region (*mxan18-19*) as indicated.

*E. coli* strains were grown in LB or 2xYT medium in the presence of relevant antibiotics or on LB plates containing 1.5% agar (Sambrook and Russel, 2001). Plasmids were propagated in *E. coli* NEB Turbo cells (New England Biolabs) (F' *proA*$^+$*B*$^+$ *lacI*$^q$ Δ*lacZM15/fhuA2* Δ*(lac-proAB) glnV galK16 galE15 R(zgb-210::Tn10)* Tet$^S$ *endA1 thi-1* Δ*(hsdS-mcrB)5*) or *E. coli* OneShot TOP10 (Invitrogen, Thermo Fisher Scientific, Waltham, USA) (F-*mcrA* Δ(*mrr-hsd*RMS-*mcr*BC) Φ80*Lac*ZΔM15 Δ *Lac*X74 *rec*A1 *ara*D139 Δ(*araleu*) 7697 *gal*U *gal*K *rps*L (StrR) *end*A1 *nup*G). Growth of *E. coli* was measured as an increase in OD at 600 nm.

Plasmid constructions. For the generation of pBR plasmids, they were propagated in *E. coli* OneShot TOP10 (Invitrogen, Thermo Fisher Scientific, Waltham, USA). Seamless assembly was used to introduce larger fragments into plasmid backbones. DNA fragments and vector backbones were amplified by PCR with primers that contained 15-20 bp overlaps between adjacent fragments. The PCR products were combined using GeneArt Seamless Cloning and Assembly Enzyme Mix (Thermo Fisher Scientific, Waltham, USA) according to manufacturer's instruction. All pDS, pAH, pKA and pPK plasmids were propagated in *E. coli* NEB Turbo cells (New England Biolabs). These plasmids were generated using standard restriction enzymes and ligated using T4-Ligase (New England Biolabs). All DNA fragments, generated by PCR were verified by sequencing.

For the introduction of point mutations or small peptide sequences blunt end cloning was employed. The entire vector was amplified with two primers extended by the sequence to be introduced. After PCR the product was digested with DpnI and the blunt ends of the PCR products were phosphorylated using T4 Phosphokinase and subsequently ligated with T4 DNA Ligase (Thermo Fisher Scientific).

For pDS79 *mCh* was amplified from pKA28 with mCherry fwd Start XbaI and mCherry stop rev HindIII and cloned into pKA45 with XbaI and HindIII.

For pDS132 *pomY*$^{CC\text{-}HEAT}$*-mCh* was amplified from pSWU30-*pomY*$^{CC\text{-}HEAT}$*-mCh* with MXAN_0634-6 and mCherry stop rev HindIII and cloned into pKA45 with XbaI and HindIII.

For pSWU30-*pomY*$^{CC\text{-}HEAT}$-*mCh* *pomY*$^{CC\text{-}HEAT}$ was amplified from genomic DNA with DS1 and DS2 and cloned into pKA28 with EcoRI and BamHI.

For pDS331 *pomY-mCh* was amplified from pDS7 with DS14 and DS40 and cloned into pMR3690 with NdeI and EcoRI.

For pAH14 *mCh* was amplified from pKA28 with KA477 and KA478 and cloned into pET45b+ with BamHI and HindIII.

For pAH180 Cys-*ftsZ* was amplified from genomic DNA with AH159 and KA500 and cloned into pET24b+ with NdeI and BamHI.

For pAH187 *pomY*$^{CC\text{-}HEAT}$-*mCh* was amplified from pDS132 with NdeI PomY fwd and DS274 and cloned into pET24b+ with NdeI and HindIII.

For pAH194 *pomY-mCh* was amplified from pDS8 with DS14 and DS274 and cloned into pET24b+ with NdeI and HindIII.

For pAH195 *pomY*$^{HEAT\text{-}IDR}$-*mCh* was amplified from pDS7 with DS12 and DS274 and cloned into pET24b+ with NdeI and HindIII.

For pAH196 *pomY*$^{CC}$-*mCh* was amplified from pPK29 with DS14 and DS274 and cloned into pET24b+ with NdeI and HindIII.

For pAH197 *pomY*$^{HEAT}$-*mCh* was amplified from pPK45 with DS12 and DS274 and cloned into pET24b+ with NdeI and HindIII.

For pAH198 *pomY*$^{IDR}$-*mCh* was amplified from pDS7 with DS271 and DS274 and cloned into pET24b+ with NdeI and HindIII.

For pAH205 *pomX-Strep* was amplified from gen. DNA DK1622 with DS278 and DS276 and cloned into pRSF-Duet1 with NcoI and HindIII.

For pAH225 *mCh* was amplified from pDS7 with mCherry fwd NdeI and DS40 and cloned into pMR3690 with NdeI and EcoRI.

For pPK29 *pomY*$^{CC}$ was amplified from pDS7 with DS14 and CC(nat) linker rev BglII and cloned into pKA45 with NdeI and BamHI.

For pPK30 *pomY*$^{HEAT}$ was amplified from pDS7 with 17 HEAT(nat) XbaI fwd and 18 HEAT(nat) linker rev BglII and cloned into pKA45 with XbaI and BglII. For cloning pKA45 was opened with XbaI and BamHI.

For pPK36 *pomY*$^{CC\text{-}HEAT}$ was amplified from pDS7 with the primers DS14 and 18 HEAT(nat) linker rev BglII and cloned into pKA28 with NdeI and BamHI. The obtained plasmid was used as a template for the following PCR with the primers DS4 and mCherry stop rev HindIII and cloned into pKA45 with XbaI and HindIII.

For pPK45 *pomY$^{HEAT}$-mCh* was amplified from pPK30 with the primers 27 HEAT fwd and mCherry stop rev HindIII and clone into pDS7 with XbaI and HindIII.

For pPK47 *pomY$^{HEAT\text{-}IDR}$-mCh* was amplified from pDS7 with the primers 27 HEAT fwd and mCherry stop rev HindIII and cloned into pDS7 with XbaI and HindIII.

For pPK46 *pomY$^{IDR}$-mCh* was amplified from pDS7 with the primers FM4 and mCherry stop rev HindIII and cloned into pDS7 with XbaI and HindIII.

pBR35_pET45b_His6-PomZ-mCherry was generated by seamless assembly. The backbone was amplified by PCR with primers BR68 and BR70 from pKA3, and the insert was amplified from pDS43 with primers BR64 and BR69.

pBR36_pRSF-Duet-1_$His_6$-mCh-PomX encodes *$His_6$-mCh-pomX* and was amplified by PCR from the plasmid pDS37 with the primers BR71 and BR72 and subsequently ligated using blunt end cloning.

pBR38_pET24b_PomY-YFP-His6 was generated by seamless assembly. The backbone was amplified by PCR with primers BR62 and BR75 from pDS3, and the fragment encoding *pomY-yfp* was amplified from pDS46 with primers BR64 and BR77.

pBR39_pET24b- PomX-His-GS-Cys encodes *pomX-$His_6$-Cys* and was amplified by PCR from the plasmid pEMR3 with the primers BR87 and BR153 and subsequently ligated using blunt end cloning.

pBR72_pET24_PomY-mCherry-His6 encodes *pomY-mCh-$His_6$.* This plasmid was generated by seamless assembly. The backbone was amplified from pBR38_pET24b_PomY-YFP-$His_6$ with primers BR62 and BR185, the insert encoding *mCh* was amplified from pBR35_pET45b_His6-PomZ-mCherry with primers BR64 and BR184.

pDS121, and pDS123. For both plasmids *pomY* was amplified from genomic DNA using primers Mxan0634 fwd BACTH and Mxan_0634-17 pomY-rev. The resulting PCR fragment was digested with KpnI and XbaI and cloned into pKT25 and pKNT25.

pDS261, pDS262, pDS263, and pDS264. For all four plasmids *pomY$^{CC}$* was amplified from genomic DNA using the primers Mxan0634 fwd BACTH and Mxan0634 Coil KpnI BACTH. The resulting PCR fragment was digested with KpnI and XbaI and cloned into pUT18, pUT18C, pKT25, and pKNT25 using the same restrictions sites.

pDS265, pDS266, pDS267, and pDS268. For all four plasmids *pomY$^{HEAT}$* was amplified from genomic DNA using the primers Mxan0634 HEAT XbaI BACTH and Mxan0634 HEAT KpnI BACTH. The resulting PCR fragment was digested with KpnI and XbaI and cloned into pUT18, pUT18C, pKT25, and pKNT25 using the same restrictions sites.

pDS269, pDS270, pDS271, and pDS272. For all four plasmids *pomY$^{IDR}$* was amplified from genomic DNA using the primers Mxan0634 Pro XbaI BACTH and Mxan_0634-17 pomY-rev. The resulting PCR fragment was digested with KpnI and XbaI and cloned into pUT18, pUT18C, pKT25, and pKNT25 using the same restrictions sites.

Fluorescence microscopy and live-cell imaging. Fluorescence microscopy was performed as described (Schumacher et al., 2017). Briefly, exponentially growing cells were transferred to slides with a thin 1.0% agarose pad (SeaKem LE agarose, Cambrex) with TPM buffer (10 mM Tris-HCl pH 7.6, 1 mM $KH_2PO_4$ pH 7.6, 8 mM $MgSO_4$), covered with a coverslip and imaged using a temperature-controlled Leica DMi6000B inverted microscope with a 100x HCX PL FLUOTAR objective at 32°C. Phase-contrast and fluorescence images were recorded with a Hamamatsu ORCA-flash 4.0 sCMOS camera using the LASX software (Leica Microsystems). Time-lapse microscopy was performed as described (Schumacher and Søgaard-Andersen, 2018). Briefly, cells were transferred to a coverslip mounted on a metallic microscopy slide and covered with a pre-warmed 1% agarose pad supplemented with 0.2% casitone in TPM buffer. Slides were covered with parafilm to retain the humidity of the agarose, and live-cell imaging was performed at 32°C. Image processing was performed with Metamorph v 7.5 (Molecular Devices). For image analysis, cellular outlines were obtained from phase-contrast images using Oufti and manually corrected if necessary (Paintdakhi et al., 2016). Fluorescence microscopy image analysis was performed with a custom-made Matlab script (Matlab R2018a, MathWorks) as described (Schumacher et al., 2021).

Structured illumination microscopy (SIM). SIM was performed on a temperature-controlled AxioObserver with Zeiss Elyra 7 Lattice SIM module with perfect focus and an α Plan-Apochromat 63x/1.46 oil Korr M27. Images were recorded with a pco.edge 4.2 sCMOS camera (PCO). Cells were mounted onto a 1% agarose pad as described above. mCh-PomX and PomY-mCh images were acquired using a 561nm 500mW laser with 100ms exposure time and 1.5% or 3% laser power, respectively. SIM images were reconstructed from 13 frames without rescaling using Zen software (Zeiss). Image processing was performed as described above.

*In vivo* fluorescence recovery after photobleaching (FRAP). *In vivo* FRAP experiments were performed as previously described (Schumacher et al., 2017) with a temperature-controlled Nikon Ti-E microscope with Perfect Focus System and a CFI PL APO 100x/1.45 Lambda oil objective at 32°C with a Hamamatsu Orca Flash 4.0 camera using NIS Elements AR 2.30 software (Nikon). Photobleaching of PomY-mCh (SA7000) was performed with two laser pulses with 30% laser power (561nm) and a dwelling time of 500msec. For mCh-PomX signals (strains SA4797 and SA9734), bleaching was performed with two laser pulses with 35% laser power (561nm) using the same dwelling time. For every image, total integrated cellular fluorescence in a region of interest (ROI) within the outline of the cell was measured together with the total integrated background fluorescence of an ROI of the same size placed outside of the cell. Additionally, fluorescence intensity was measured in the bleached region together with background fluorescence of an ROI of the same size placed on the background. After background correction, the corrected fluorescence intensity of the bleached ROI was divided by total corrected cellular fluorescence, which corrects for bleaching effects during picture acquisition. This relative fluorescence was correlated to the initial fluorescence in the bleached ROI. The mean relative fluorescence of several cells was plotted as a function of time [min]. The recovery rate for the tested fluorescent protein (t1) was determined by fitting the plotted data to a single exponential equation (y=y0+A*e-x/t) using Graphpad Prism 9.0.2 (161). Half-maximal recovery (t1/2) was calculated from the recovery rate (t1) by t1/2= ln (2) * t1.

Bacterial two hybrid assays (BACTH). BACTH experiments were performed as described (Karimova et al., 1998). Relevant genes were cloned into the appropriate vectors to construct N-terminal and C-terminal fusions with the 25 kDa N-terminal or the 18 kDa C-terminal adenylate cyclase fragments of *Bordetelly pertussis*. Restoration of cAMP production was observed by the formation of blue color on LB agar supplemented with 80 µg/ml 5-bromo-4-chloro-3-indolyl-β-D-galactopyranoside (X-Gal) and 0.25 mM isopropyl-β-D-thiogalactoside (IPTG). As positive control, the leucine zipper from GCN4 was fused to the T18 and the T25 fragment. As a negative control, plasmids were co-transformed that only expressed the T18 or T25 fragment. All tested interactions were spotted on the same LB agar plates with positive control and all corresponding negative controls. Proteins were tested for their interaction in *E.coli* BTH101 **(**F- *cya-99 araD139 galE15 galK16 rpsL1* (StrR) *hsdR2 mcrA1 mcrB1* cells.

Immunoblot analysis. Samples were prepared by harvesting exponentially growing *M. xanthus* cells and subsequent resuspension in SDS lysis buffer to an equal concentration of cells. Western blot analyses were performed as described before (Sambrook and Russel, 2001) with rabbit polyclonal α-PomX (1:10000), α-PomY (1:10000) (Schumacher et al., 2017), α-FtsZ (1:10000) (Treuner-Lange et al., 2013), α-PilC (1:3000) (Bulyha et al., 2009), α-GFP (1:2500, Roche) or α-mCh (1:10000; Biovision) primary antibodies together with either horseradish-conjugated goat α-rabbit immunoglobulin G (1:25000) (Sigma-Aldrich) or horseradish-conjugated sheep α-mouse immunoglobulin G (1:2000) (Amersham) as the secondary antibody (applies only for α-GFP western blots). For PomY blots, protein transfer was performed with a Tris/CAPS buffer system (BioRad). Protein transfer was otherwise accomplished with Trans-Blot Turbo buffer (BioRad) for all other blots. Blots were developed using Luminata Forte Western HRP Substrate (Millipore) and visualized using a LAS-4000 luminescent image analyzer (Fujifilm).

Protein purification. PomY-His$_6$ and native FtsZ were purified as described (Schumacher et al., 2017). Cys-FtsZ, has an N-terminal MGKCKGSG extension than can be labelled with a fluorophore after purification. Cys-FtsZ was purified as native FtsZ. To purify PomY-mCh-His$_6$ (analogous to the active PomY-mCh fusion *in vivo*) plasmid pAH194 was propagated in *E. coli* ArcticExpress(DE3) RP cells (Agilent Technologies), grown in 2xYT medium with 50µg/ml kanamycin at 30°C to an $OD_{600}$ of 0.6–0.7. Protein expression was induced with 1mM IPTG for 16h at 18°C. Cells were harvested by centrifugation at 5000×*g* for 20min at 4°C and washed in lysis buffer 1 (50mM $NaH_2PO_4$, 300mM NaCl, 10mM imidazole, 20mM β-mercaptoethanol, pH 8.0). Cells were then lysed in 50ml lysis buffer 2 (lysis buffer 1, 100µg/ml phenylmethylsulfonyl fluoride (PMSF), 1× complete protease inhibitor (Roche Diagnostics GmbH), 10U/ml benzonase, 0.1% Triton X-100, pH 8.0) by three rounds of sonication for 5min with a UP200St ultrasonic processor (pulse 80%, amplitude 80%) (Hielscher) on ice. Cell debris was removed by centrifugation (4,150× *g* for 45min at 4°C) and filtration with a 0.45µm sterile filter (Millipore Merck). PomY-mCh-His$_6$ was purified with a 5ml HiTrap Chelating HP (Cytiva, 17040901) column preloaded with $NiSO_4$ and equilibrated with lysis buffer 1. The column was washed with 20 CV (column volumes) wash buffer 1 (lysis buffer 1, 20mM imidazole) and five CV wash buffer 2 (lysis buffer 1, 50mM imidazole). Protein was eluted with elution buffer (lysis buffer 1, 300mM imidazole). Protein was then applied to a HiLoad 16/600 Superdex 200pg (Cytiva; 28989335) equilibrated in dialysis buffer (50mM HEPES/NaOH, 150mM KCl, 0.1mM EDTA, 20mM β-mercaptoethanol,

10% (v/v) glycerol, pH 7.2). Fractions with PomY-mCh-$His_6$ were snap-frozen in liquid nitrogen and stored at -80°C until use. Protein concentration was determined immediately before use using Protein Assay Dye Reagent Concentrate (BioRad). Before use, proteins were dialyzed in dialysis buffer 2 (dialysis buffer 1, 50mM KCl). $PomY^{CC}$-mCh-$His_6$, $PomY^{CC-HEAT}$-mCh-$His_6$, $PomY^{HEAT}$-mCh-$His_6$, $PomY^{HEAT-IDR}$-mCh-$His_6$, and $PomY^{IDR}$-mCh-$His_6$ were purified like PomY-mCh-$His_6$ using the plasmids pAH196, pAH187, pAH197, pAH195, and pAH198, respectively.

PomX-$His_6$, PomX-$His_6$-Cys, and $His_6$-mCh-PomX (which is similar to the active mCh-PomX fusion *in vivo*) were purified under denaturing conditions and subsequently refolded. To purify them, the plasmids pEMR3, pBR39 and pBR36, respectively, were propagated in Nico21 (DE3) (NEB). Cells were grown in LB media with 50µg/ml kanamycin at 30°C until an $OD_{600}$ of 0.5. After cooling of cells to 18°C, expression was induced with 0.4mM IPTG for 16h at 18°C. Cells were harvested by centrifugation at 4500× *g* for 10min at 4°C. Cell pellets were subsequently frozen at -80°C. For purification, cell pellets were resuspended in PomX lysis buffer (50mM $NaH_2PO_4$, 300mM NaCl, 10mM imidazole, 8M urea, pH 8.0, 1× complete protease inhibitor (EDTA free) (Roche Diagnostics GmbH), 0.4mM Tris(2-carboxyethyl)phosphine hydrochloride (TCEP), 10U/ml DNase 1, 100µg/ml lysozyme) and lysed by sonication with a tip sonicator (2.30 min, 20 s pulses, 30% amplitude). After centrifugation to clear cell debris (25,000× *g,* 45min, 4°C), the lysate was incubated with Ni-NTA agarose (Qiagen, Hilden, Germany) for 16h in a 50 ml plastic tube. Subsequently, the Ni-NTA agarose was washed six times with 45ml of PomX wash buffer (50mM $NaH_2PO_4$, 300mM NaCl, 20mM imidazole, 6M urea, 0.4mM TCEP, pH 8.0) by resuspension and centrifugation (500× *g*, 4°C, 5 min). In the last step, the Ni-NTA agarose was applied to a gravity column and eluted with PomX elution buffer (50mM $NaH_2PO_4$, 300mM NaCl, 250mM imidazole, 6M urea, 0.4mM TCEP, pH 8.0). PomX-$His_6$-Cys containing protein fractions were used for labeling before refolding. Protein containing fractions were pooled for refolding during dialysis in Slide-A-Lyzer cassette (Thermo Fisher Scientific) with a 10,000 MWCO. Refolding was achieved with four subsequent dialysis steps with decreasing amounts of urea ($1^{st}$ dialysis: 4M urea for 4h, $2^{nd}$: 2M urea for 16h, $3^{rd}$ and $4^{th}$: 0M urea for 4h each) in the PomX dialysis buffer (50mM HEPES/NaOH, 50mM KCl, 0.1mM EDTA, urea, 10% (v/v) glycerol, pH 7.2) at 4°C. Proteins were frozen in liquid nitrogen and stored at -80°C until further use.

For purification of PomX-Strep, plasmid pAH205 was propagated in *E. coli* Rosetta2(DE3) cells, grown in 2xYT medium with 50µg/ml kanamycin, 30µg/ml chloramphenicol, and 0.5% glucose at 37°C to an $OD_{600}$ of 0.6-0.7. Protein expression was induced with 0.5mM IPTG for 18h at 18°C. Cells were harvested and washed in StrepTag lysis buffer 1 (100mM Tris-HCl pH 8.0, 150mM NaCl, 1mM EDTA, 1mM β-mercaptoethanol) and lysed in StrepTag lysis buffer 2 (StrepTag lysis buffer 1, 100µg/ml PMSF, 1× complete protease inhibitor (Roche Diagnostics GmbH), 10U/ml benzonase) by four rounds of sonication for 5min with a UP200St ultrasonic processor (pulse 80%, amplitude 80%) (Hielscher) on ice. Cell debris was removed by centrifugation (4,150× *g* for 45min at 4°C). PomX-Strep was purified from a batch using 5ml Strep-TactinXT 4Flow resin (iba), equilibrated with StrepTag lysis buffer 1. The resin was incubated with the cleared lysate for 18h at 4°C on a rotary shaker. Contaminating proteins were eluted from the resin by washing 5× with 10ml StrepTag lysis buffer 1. The protein was eluted with 1× BXT buffer (100mM Tris-HCl pH 8.0, 150mM NaCl, 1mM EDTA, 50mM biotin) (iba). Fractions containing PomX-Strep were pooled and dialyzed against 3× 5l of dialysis buffer at 4°C. Proteins were frozen in liquid nitrogen and stored at -80°C until used.

Protein labeling. For labeling of PomX-$His_6$-Cys the protein containing fractions were mixed with 125µg Alexa Fluor 488 $C_5$ Maleimide (Thermo Fisher Scientific) and incubated for 2h at RT under denaturing conditions. Subsequently, unreacted dye was removed by repeated concentration and dilution in Amicon Ultra 100 kDa MWCO filters (Merck Millipore, Darmstadt, Germany) using dialysis buffer (50mM HEPES/NaOH, 50mM KCl, 0.1mM EDTA, 6M urea, 2.5% (v/v) glycerol, pH 7.2) at 4°C. Afterwards the protein was refolded via dialysis as described above. The labelled protein is indicated as PomX-A488. Cys-FtsZ contains an N-terminal MGKCKGSG extension to FtsZ. For labeling of Cys-FtsZ the protein was mixed with 125µg Alexa Fluor 488 $C_5$ Maleimide (Thermo Fisher Scientific) for 2h at 4°C before unreacted dye was separated from the protein using an Econo-Pac 10DG desalting column (Biorad, Hercules, USA). The labelled protein is indicated as A488-FtsZ.

FtsZ GTPase assay. GTP hydrolysis by FtsZ was determined using a 96-well NADH-coupled enzymatic assay with modifications. Assays were performed in reaction buffer (50mM HEPES/NaOH pH 7.2, 50mM KCl, 10mM $MgCl_2$) with 0.5mM nicotinamide adenine dinucleotide (NADH) and 2mM phosphoenolpyruvate and 3µl of a pyruvate kinase/lactate dehydrogenase mix (PYK/LDH; Sigma). Buffer was pre-mixed with proteins in low-binding microtubes (Sarstedt) on ice. If necessary, a dialysis buffer was added to correct for glycerol in the assays. 100µl mixtures were transferred into transparent UV-STAR µCLEAR 96-well microplates (Greiner bio-one). The addition of 1mM GTP started the reaction. Measurements were performed in an infinite M200PRO (Tecan) for 2h in 30s intervals at 32°C, shaking at 340 nm wavelength. Assays were also performed without the addition of FtsZ or Cys-FtsZ, and these measurements were subtracted to account for background by spontaneous GTP hydrolysis and UV-induced NADH decomposition. The light path was determined to be 0.248cm with known NADH concentrations. The extinction coefficient of NADH ε340 = 6220 $M^{-1}cm^{-1}$ was used. Experiments were replicated at least three times with independent protein preparations.

FtsZ sedimentation assay. Sedimentation assays were performed as described (Schumacher et al., 2021). Briefly, before sedimentation experiments, FtsZ and Cys-FtsZ were applied to a clear spin at 20,000× *g* for 10min at 4°C. Proteins at a final concentration of 10µM in a total volume of 50µl were pre-mixed and incubated for 5min at 32°C in a buffer (50mM Hepes/NaOH pH 7.2, 50mM KCl, 10mM $MgCl_2$, 10mM $CaCl_2$, 1mM β-mercaptoethanol) and after 10min GTP was added to 5 mM. Samples were separated into soluble and pellet fractions by high-speed centrifugation at 160,000× *g* for 1h at 25°C. Soluble and pellet fractions were separated, and volumes were adjusted with 1×SDS sample buffer. Fractions were separated by SDS-PAGE and stained with Instant Blue (expedion) for 10min. Experiments were replicated at least three times with independent protein preparations.

*In vitro* pull-down experiments. 10µM protein alone or pre-mixed as indicated were incubated for 1h at 32°C in reaction buffer (50mM HEPES/NaOH pH 7.2, 50mM KCl, 10mM $MgCl_2$) in a total volume of 200µl and applied to 20µl 5% (v/v) MagStrepXT beads (iba) for 30min. Magnetic beads were washed 10× with 200µl reaction buffer. Proteins were eluted with

200µl 1× BXT buffer (100mM Tris-HCl pH 8.0, 150mM NaCl, 1mM EDTA, 50mM biotin) (iba).

Turbidity measurements. Turbidity measurements were set up using a liquid handler (Freedom EVO platform, Tecan) in flat bottom UV-STAR 384 well plates (#781801, Greiner BioOne, Frickenhausen, Germany). For assays, reaction buffer, storage buffer for adjustment, and artificial crowding agent were added to individual wells and mixed, before the addition of the proteins after which the mixture was mixed again by pipetting. All reactions were set up in triplicates on the plate. Protein was thawed immediately before addition and the protein and the 384 well plate were cooled to 4°C during the liquid handling process. All experiments were performed at a constant final buffer composition of 50mM HEPES pH 7.25, 50mM KCl, 3%Glycerol, 7mM $MgCl_2$ with or without artificial crowder (final concentration given in % w/v). Subsequently the turbidity was determined via absorption at 340 nm using a plate reader (model Spark, Tecan)

Setup of in *vitro* phase separation assays by microscopy. *In vitro* phase separation assays were either conducted in 384 well glass bottom plates (#781892, Greiner BioOne, Frickenhausen, Germany) or in sticky-Slide VI 0.4 chambers (ibidi GmbH, Gräfelfing, Germany) attached to cleaned cover slides (rinsed with ethanol and $ddH_2O$). The plates and chambers were further cleaned and hydrophilized by plasma cleaning with oxygen as the process gas (model Zepto, Diener Electronic). Directly afterwards, chambers were incubated with 25-50 µl of 0.5mg/ml PLL(20)-g[3.5]-PEG(2) (SuSoS AG, Duebendorf, Switzerland). After incubation for more than 30min at RT, chambers were washed three times with reaction buffer (50mM HEPES pH 7.2, 50mM KCl, 10mM $MgCl_2$). The concentration of crowding agent stock solution (PEG8000, Ficol 70, Ficol 400, Dextran T500) were determined by measuring the refractive indices. For assays, reaction buffer, storage buffer for adjustment, and artificial crowding agent were added to the chambers and mixed by pipetting before the addition of the proteins. Phase separation was induced by adding the proteins and gently mixing by pipetting. All experiments were performed at a constant final buffer composition of 50mM HEPES pH 7.25, 50mM KCl, 3% glycerol, 7mM $MgCl_2$ with or without artificial crowder (final concentration given in % w/v) at a constant room temperature of 23°C.

For experiments with PomX-A488, or experiments involving mixtures of several proteins, i.e. PomY and PomX or PomY and FtsZ or Streptavidin, experiments were performed in the presence of 4mM TCEP to prevent protein crosslinking.
To generate heat maps of the behaviour of PomY and PomY-mCh, and PomX, PomX-A488 and mCh-PomX in response to different protein and PEG8000 concentrations, individual chambers were pipetted in rows of equal protein concentration starting from the lowest crowder concentration. Images were acquired after 15 min of incubation starting from the highest crowder concentrations.

FRAP of PomY-mCh condensates was additionally performed in the presence of 4mM TCEP, an oxygen scavenger system (3.7U/ml pyranose oxidase, 90U/ml catalase, 0.8% glucose) (Swoboda et al., 2012) and Trolox to reduce the aging of PomY condensates. Condensates were formed under standard conditions, but particularly large condensates were selected for FRAP measurements.

For experiments with PomY-mCh and PomX-A488, 4μM PomX-A488 was added first to the chamber then 4μM PomY-mCh and subsequently the chamber was well mixed by pipetting up and down 7-10 times, before incubation for 1h before image acquisition.

Experiments for the determination of the enrichment of A488-FtsZ or Streptavidin-A488 in PomY condensates, 4μM PomY-mCh was first added to the chamber then 0.5 or 1μM A488-FtsZ or Strep-A488 and subsequently gently mixed by pipetting, before incubation for 30min before image acquisition.

Experiments showing FtsZ filament formation from PomY condensates were conducted with 2% PEG8000, 4μM PomY-mCh, 1μM A488-FtsZ, 4mM TCEP, the oxygen scavenger system and Trolox.

<u>Fluorescence microscopy of *in vitro* phase separation assays.</u> All images unless otherwise mentioned were acquired on a Zeiss LSM780 confocal laser scanning microscope using a Zeiss C-Apochromat 40x/1.20 water-immersion objective (Carl Zeiss AG, Oberkochen, Germany). Tile scans and time-series were recorded using the built in definite focus system. All two-color images were acquired with alternating illumination to avoid cross-talk. FtsZ-A488/PomX-A488 were excited using the 488nm Argon laser, PomY-mCh or mCh-PomX constructs using the 561nm DPSS laser. Images were usually acquired as a tile-scan and confocal Z-stack with about 10-50 planes and optimized slicing. Images for analysis were typically recorded with a pinhole size of 14 Airy units, 512×512 pixel resolution, and a scan rate of 1.27μs per pixel. Images for display were typically recorded with the same settings, but also at a scan zoom of 4 and a line averaging of 4. Images to visualize FtsZ filament formation from PomY-mCh condensates were typically recorded with a pinhole size of 1 Airy unit, 512×512 pixel resolution, and a scan rate of 1.27μs per pixel. Time-series to observe the fusion of PomY-mCh condensates were typically recorded with ~1 s intervals, while time-series of FtsZ filament formation originating from PomY-mCh condensates were typically acquired with ~30s intervals.

For FRAP experiments photobleaching of large PomY-mCh condensates (diameter: ~ 9-15μm in) was performed with ten laser pulses with 100% laser power (561nm). The region of interest (ROI) selected for FRAP was centered in the middle of the condensates (diameter: ~3.5μm) and time-series to record fluorescent recovery were acquired at ~2-5s intervals. FRAP analysis was performed as in the *in vivo* FRAP.

<u>Image analysis for *in vitro* assays.</u> All images were processed using Fiji (Schindelin et al., 2012) (version v1.53q). Images were analyzed using custom-written Fiji macros. The phase separation of PomY-mCh under various conditions was quantified by estimating the separation factor. The condensates in each image were identified and segmented from the maximum intensity projection of individual z-stacks, by means of the StarDist plugin, using the 2D pretrained model *2D_versatile_fluo* (Schmidt et al., 2018). The segmented condensates were then screened, and those with an intensity below three standard deviations of the background signal or a size of less than nine pixels (corresponding to a diameter of 300nm) were removed. Intensity values inside and outside the condensates were measured separately, and used to calculate the separation factor, in percentage, as the integrated intensity of the condensates over the total integrated intensity of the image which reflects the total amount of protein that is located in the condensed phase.

To quantify the recruitment of proteins into PomY-mCh condensates we used a similar approach, with the following differences. Instead of using the maximum intensity projection of the images, the slice that was best in focus, i.e. with the maximum standard deviation of the intensity, was selected for analysis of enrichment and separation rate. The PomY-mCh channel was used to determine location and size of the condensates, as before. The segmentation was used to estimate the fluorescence intensity in the condensates as well as in the background for all three signals of interest, PomY-mCh, A488-FtsZ and Strep-A488. Besides estimating the separation factor in PomY-mCh, the enrichment of A488-FtsZ and Strep-A488 in the condensates was computed as the ratio between the average intensity of the intensity in all the condensates and the average intensity in the remaining areas of the field of view, that is the background.

Statistics. The mean and standard deviation (STDEV) were calculated with Graphpad Prism 9.0.2 (161). Localization patterns from fluorescence microscopy data, cell length distribution, and constrictions frequency were quantified based on the indicated number of cells per strain in three independent experiments unless otherwise indicated. Scatter dot plots were generated with GraphPad Prism 9.0.2 (161). Statistical analysis was performed with GraphPad Prism 9.0.2 (161). All data sets were tested for significant differences using a 2way ANOVA with multiple comparisons. A *P*-value $<0.01$ was used to determine statistically significant significant difference.

Bioinformatics. PomY orthologs were identified in a best-best hit reciprocal BlastP analysis from fully-sequenced genomes of myxobacteria (Altschul et al., 1990). Protein domains were predicted using SMART (Letunic and Bork, 2018) and Interpro (Blum et al., 2021). Disorder prediction was performed with IUPred2A using the IUpred2 and Anchor2 algorithms (Erdos and Dosztanyi, 2020; Meszaros et al., 2018). Isoelectric points were predicted with the ProtParam tool using the Expasy webserver (Gasteiger et al., 2003).

Data availability. The authors declare that all data supporting this study are available within the article or its Supplementary Information file.
